# Asymmetry and niche partitioning shape the infection dynamics of co-transmitted *Wolbachia* symbionts

**DOI:** 10.64898/2026.07.08.737353

**Authors:** Megan W. Jones, Pablo A. Stilwell, Amelia R. I. Lindsey

**Affiliations:** Department of Entomology, University of Minnesota, St. Paul, Minnesota, USA

**Keywords:** symbiosis, host-microbe interaction, intracellular, *Wolbachia*, vertical transmission

## Abstract

*Wolbachia* is an incredibly widespread maternally transmitted bacterium in arthropods that can alter host physiology, nutrition, reproduction, and immunity. In some cases, multiple *Wolbachia* strains infect the same host and are stably transmitted alongside each other. This raises the question of how multiple intracellular symbionts interact with each another and with the host to ensure stable transmission. Here, we use fluorescence *in situ* hybridizations and confocal microscopy to investigate co-transmission in a naturally occurring co-infection of two *Wolbachia* strains in *Drosophila simulans*: *w*Ha and *w*No. We find significant differences in spatial occupancy and abundance between the co-transmitted strains across stages of oogenesis and embryogenesis. We show that *w*Ha and *w*No have biases for different niches during oogenesis, and their strain-specific abundance is driven by egg chamber development, mating status, and their interaction. After differential curing of the co-infection, we find that *w*No is dependent on *w*Ha for vertical transmission, but not vice versa. Additionally, while *w*Ha localization patterns are unchanged by loss of co-infection, abundance of *w*Ha in the ovaries increases when *w*No is removed. Understanding how symbiont co-infections achieve stability has important implications for the ongoing use of *Wolbachia* as a tool for insect management programs, but also for our understanding of the ecology of intracellular communities more broadly.

## INTRODUCTION

Intracellular microbes are taxonomically and functionally diverse among arthropods. Many are vertically transmitted and induce a wide variety of host effects that favor symbiont transmission. For example, intracellular microbes can modify host metabolism and regulate development [1–3], supplement essential nutrients [4–9], alter host reproduction [10, 11], or provide defense from pathogens and parasitoids [12–17]. Symbiont-mediated impacts are not mutually exclusive and symbionts are often capable of impacting myriad host processes, both within an individual host and sometimes across multiple host species [18]. A single host may also harbor multiple symbiont strains or species, adding an additional level of complexity to symbiont-host dynamics. Indeed, many arthropods have two or more co-transmitted intracellular symbionts [19] and some notable examples contain three, four, or even five co-transmitted intracellular symbionts [19–23]. Despite how common co-infections are among arthropods, we understand very little about how these relationships are formed and regulated.

The most common intracellular infection in arthropods is with *Wolbachia* (Alphaproteobacteria: Rickettsiales). *Wolbachia* is primarily vertically transmitted and infects the cells of many invertebrates, including plant-parasitic and filarial nematodes, terrestrial crustaceans (Isopoda), non-insect hexapods (Collembola), arachnids, and many insects [24–28]. Over evolutionary timescales, *Wolbachia* are also transmitted horizontally, and have undergone numerous host switches, especially within the Arthropoda [29, 30]. *Wolbachia* infects an estimated 1/3 to 2/3 of all insect taxa, and often co-occurs with other vertically transmitted intracellular symbionts, including other strains of *Wolbachia* [19, 20, 25, 31]. In many cases, co-infections are persistent, and stably co-transmitted across host generations [32–34]. In theory, co-infecting symbionts could compete or interfere with one another, leading to host fitness costs or unstable transmission [35, 36]. Alternatively, co-infections could produce additive or complementary effects that favor co-infected hosts [33, 37–40].

It is unclear which factors facilitate stable co-infection and co-transmission but likely a combination of the host and symbionts is responsible [23, 40, 41]. We hypothesize that divergent tissue tropisms between co-infecting symbionts is one important factor that could facilitate stability. Arthropod-infecting symbionts vary significantly in the tissues they preferentially occupy, and often exhibit tropisms for specific host structures, whole organ systems, tissues, or cell types [32, 42–44]. For example, in whiteflies (*Bemisia tabaci,* Hemiptera: Aleyrodidae) harboring co-infections of *Portiera* and *Hamiltonella*, symbionts occupy the same cell types, but are spatially partitioned within the intracellular environment by endomembrane systems [45]. In the tsetse fly (Diptera: Glossinidae), co-infecting symbionts *Wolbachia*, *Wigglesworthia*, and *Sodalis* are partitioned across distinct host tissue and organ niches [32]. These patterns suggest that occupying different spatial niches may be important for mitigating competition and conferring transmission stability across generations. In *Drosophila*, there is evidence that host processes such as autophagy can restrict symbiont localization and proliferation [46], as well as evidence that symbiont niche tropism is symbiont-driven [42], suggesting that a combination of host and symbiont factors play a role. However, disentangling the interactions between host and symbiont factors that facilitate stable transmission of co-infecting symbionts is challenging, given the taxonomic diversity present in many arthropod co-infections, and a lack of tractable laboratory models.

To investigate the dynamics of stable co-transmission, we leveraged a tractable model of co-infection: two *Wolbachia* strains (“*w*Ha” and “*w*No”) that co-inhabit some *Drosophila simulans* (Diptera: Drosophilidae) [33, 40]. In nature, populations with the *w*Ha and *w*No co-infection are fixed for both strains, which are co-transmitted with high fidelity [40, 47]. Importantly, co-infected flies have the reproductive advantage over monoinfected or uninfected flies due to the ability of both *w*Ha and *w*No to induce a conditional sperm-egg lethality known as cytoplasmic incompatibility (CI) [40, 47]. Previous work in this system identified distinct patterns in the relative abundance of each *Wolbachia* strain in ovaries as compared to very young embryos, based on qPCR [48]. *w*No, which was the dominant strain in the ovaries, was present at roughly the same abundance as *w*Ha in embryos [48]. These findings suggest that strain-specific changes in infection dynamics occur during oogenesis and transmission to the next generation [48]. Additionally, there were striking differences in the relative abundance of the two *Wolbachia* strains between somatic and ovary tissues. While *w*No was the dominant strain in the ovaries, *w*Ha was dominant in the soma, hinting at strain-specific differences in niche occupancy [48]. Here, we build on these findings, to investigate co-transmission of strains through the ovaries and determine (i) localization patterns of each strain in the ovaries and whether spatial partitioning occurs during oogenesis, (ii) abundance of each strain in the ovaries during oogenesis and whether strain titers are differentially influenced by host mating, (iii) whether strain localization patterns differ between ovaries and embryos, and (iv) whether *w*Ha and *w*No localization and abundance patterns change under co-infection versus single infection conditions.

## MATERIALS AND METHODS

### Fly stocks

*Drosophila simulans* (Cornell Stock Center SKU: 14021-0251.198) were reared on Bloomington *Drosophila* Stock Center (BDSC) food (Nutri-Fly Bloomington formulation) including the recommended concentration of propionic acid. Stocks were maintained at 25°C on a 24 hour, 12:12 light-dark cycle. *Wolbachia*-uninfected stocks were previously generated [48] by treating flies with tetracycline and restoring their gut microbiome following standard protocols [49]. Stocks exposed to antibiotics were maintained for at least 12 generations without antibiotics prior to use in experiments. To generate monoinfected stocks, we used a partial heat cure to destabilize *Wolbachia* transmission, as described previously [48]. In brief, adult flies were held at 30°C for three days, then transferred to fresh food and allowed to oviposit for three days under standard rearing conditions. Female offspring of heat-treated mothers were collected upon eclosion, mated with *Wolbachia*-uninfected males (to avoid CI), and allowed to initiate isofemale lines. Once offspring were visible, the mothers were removed, flash frozen in liquid nitrogen, and stored at -20°C for *Wolbachia* screening (see below).

### *Wolbachia* screening and sequence validation

For routine screening of all stocks for *Wolbachia* infection, and for identifying isofemale lines with monoinfections, we used a multiplex PCR which targets the *Wolbachia*-specific gene *wsp* and produces size-diagnostic amplicons for *w*Ha and *w*No (Table 1) [40]. DNA was extracted from whole flies with the DNEasy Blood and Tissue kit (Qiagen). PCRs were performed in 20 μL reactions with Q5 HotStart High-Fidelity 2X master mix (New England Biolabs) with 500 nM 81F, 500 nM 635R, and 75 nM 463R primers, plus 1 µL whole fly DNA extraction as template. Samples underwent initial denaturation for two minutes at 98°C followed by 35 cycles of 10 seconds at 98°C, 15 seconds at 57°C, and 15 seconds at 72°C, and then a final extension for two minutes at 72°C. PCR products were run on 1% agarose gels, stained post-electrophoresis with GelRed (Biotium), and visualized under UV light.

**Table 1.**
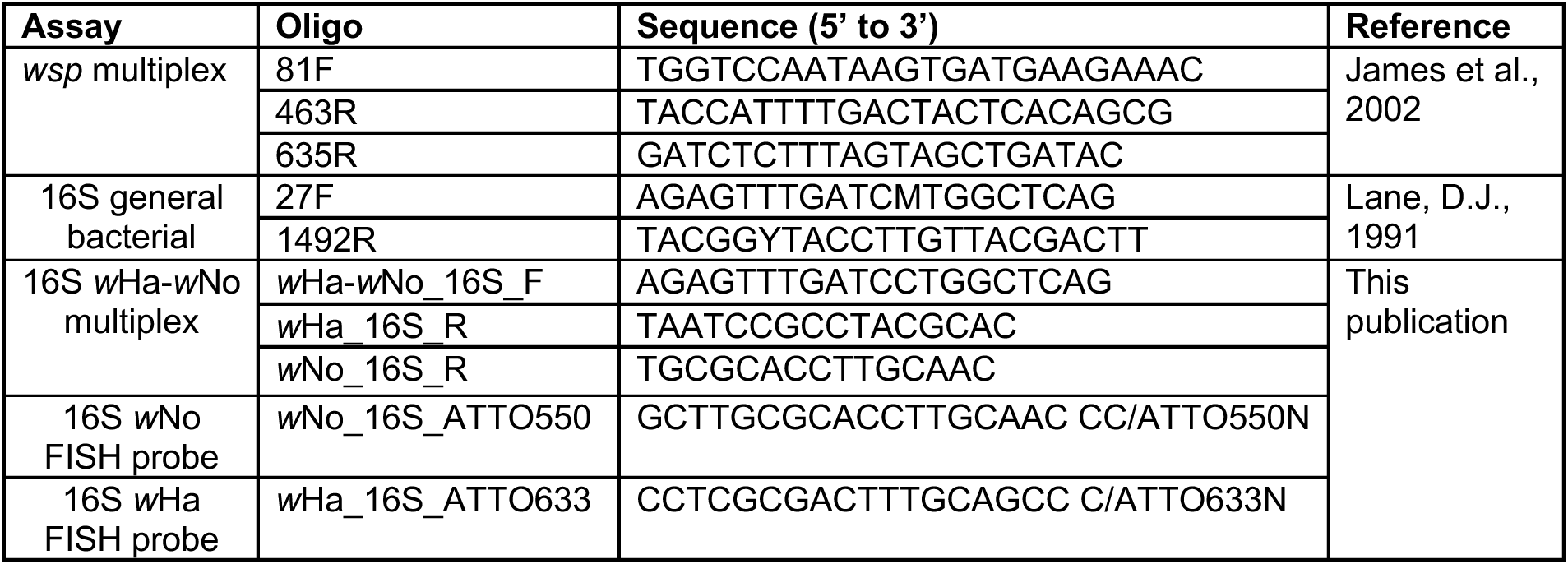
Oligonucleotides used in this publication.

### FISH probe design

We selected 16S rRNA as a fluorescence *in situ* hybridization (FISH) probe target. Prior to designing probes, we cloned and sequenced *w*Ha and *w*No 16S rRNA genes from the co-infected stock to determine if there were differences as compared to published reference genomes. DNA was extracted from dissected ovaries (to limit contamination from the gut microbiome) with the DNEasy Blood and Tissue kit (Qiagen). 16S rRNA genes were amplified with 27F/1492R general bacterial 16S primers (Table 1) in 20 µL reactions with Q5 High-Fidelity 2X master mix (New England Biolabs), 500 nM of each primer, and 1 µL ovary DNA extraction as template [50]. Samples underwent initial denaturation for two minutes at 95°C followed by 30 cycles of 15 seconds at 95°C, 15 seconds at 55°C, and 90 seconds at 72°C, followed by a final extension of two minutes at 72°C. PCR products were ligated into plasmids with the TOPO blunt end cloning kit (ThermoFisher Scientific) and transformed into One Shot TOP10 competent cells (Invitrogen). Cells were plated on Lysogeny Broth with agar and 50 µg/mL kanamycin to select for transformants and incubated at 37°C overnight. Individual colonies were subcultured and grown in Lysogeny Broth with 50 µg/mL kanamycin with shaking at 37°C overnight, and plasmids were extracted with the Monarch plasmid miniprep kit (New England Biolabs). We designed multiplex PCR primers targeting 16S that produce size-diagnostic amplicons for *w*Ha and *w*No (Table 1) to screen minipreps prior to sequencing. PCR and gel electrophoresis followed the same protocol as used for full length 16S PCRs (see above), instead with a 62°C annealing temperature and 40 second extension time during cycling. Six plasmid minipreps (three *w*Ha 16S, three *w*No 16S) were submitted for Sanger sequencing of the inserts in both directions. Consensus sequences were generated with SnapGene and aligned against published *w*Ha (NC_021089.1) and *w*No (NC_021084.1) 16S rRNA sequences with MAFFT [51, 52]. Following sequence validation, probes were selected as ∼20bp regions where the two strains differed significantly (≥3 bp) but had similar GC content (Table 1). Probes were mapped against *w*Ha (NC_021089.1), *w*No (NC_021084.1), and *Drosophila simulans* (GCF_016746395.2) reference genomes to check for off-target alignments. 3’ ATTO 550 and ATTO 633 fluorescent tags (ATTO-TEC) were selected to minimize spectral overlap with each another and with host tissue autofluorescence.

### FISH: ovary collection and fixation

Adult females were isolated upon emergence in vials with BDSC food. To generate unmated females, isolated females were allowed to mature for three days in the absence of males. To generate mated females, isolated females were allowed to mature for two days then provided with an equivalent number of *Wolbachia*-uninfected males one day prior to dissection. One day prior to sample collection, all flies were provided with yeast paste to promote egg development. Ovary fixation protocols were adapted from a published FISH protocol for *Wolbachia* detection in *Drosophila* ovaries [53]. FISH sample processing was performed under RNAse-free conditions as follows. Age-matched flies of different infection statuses were dissected in ice-cold diethyl pyrocarbonate-treated phosphate buffered saline solution (DEPC-PBS) then fixed in a 50:50 mixture of 4% paraformaldehyde and heptane, with nutation for 30 minutes. Samples were washed 3 times with 500 μL 50:50 methanol:DEPC-PBS, then 3 times in 100% methanol. Samples were stored in 100% methanol at -20°C.

### FISH: embryo collection and fixation

To collect embryos, 30-50 adult *Drosophila simulans* (aged <7 days) were placed in mating cages with grape agar plates (3% agar, 25% storebought grape juice, 0.3% sucrose) streaked with yeast paste. Embryos were washed off plates with sterile PBS and collected in 70 µm cell strainers. Embryos underwent additional processing and permeabilization steps as compared to ovaries, adapted from Rothwell and Sullivan [54]. First, embryos were dechorionated for 3 minutes in 50% bleach then rinsed in DEPC-PBS. Dechorionated embryos were transferred to microcentrifuge tubes with a 50:50 mixture of heptane and methanol, repeatedly struck hard against the bench for 1 minute, and then allowed to settle. This step was repeated for a total of 3 times. Methanol, heptane, and embryos remaining at the liquid interface were removed. Samples were washed 3 times with 500 μL 50:50 methanol:DEPC-PBS, followed by 3 times in 100% methanol. Samples were stored in 100% methanol at -20°C prior to staining.

### FISH staining

Methanol from storage was removed and samples were rehydrated stepwise in increasing fixative concentrations (percentage methanol: percentage 4% paraformaldehyde in DEPC-PBS: 5 m 70:30, 5 m 30:70, 20 m 0:100). Samples were washed 3 times in 500 μL DEPC-PBS with 1% Tween-20 (DEPC-PBST) before incubating in 500 μL DEPC-PBST with 50 ng/μL proteinase K for 10 minutes with shaking (350 rpm) at 37°C. Following proteinase K treatment, samples were fixed a final time in 4% paraformaldehyde in DEPC-PBS for 20 minutes with nutation, then washed three times with DEPC-PBST. Samples were incubated in 500 μL of a 50:50 mix of DEPC-PBS and hybridization buffer (50% formamide, 1X saline sodium citrate buffer (SSC), 0.5X Denhardt’s solution (Thermo Fisher), 0.5% ssDNA (Invitrogen), 50 mM TrisHCl, 0.1% SDS) at 42°C for 5 minutes. Buffer was removed, and samples were incubated in 100% hybridization buffer at 42°C for 5 minutes, followed by a second incubation in fresh 100% hybridization buffer at 42°C for 30 minutes. All subsequent steps were performed in the dark. Probes were added to hybridization buffer to a final concentration of 0.397 μM and samples were incubated at 42°C overnight. For no probe controls, samples were incubated at 42°C overnight in hybridization buffer only. Following overnight incubation, all samples were washed twice for 15 minutes each at 42°C with wash buffer 1 (1X SSC, 0.1% SDS, 20 mM Tris-HCl) then twice for 15 minutes each at 42°C with wash buffer 2 (0.5X SSC, 0.1% SDS, 20 mM Tris-HCl). Following washes, all samples were incubated in 1 μg/mL DAPI at 4°C overnight. Samples were arranged on slides in 25 μL Prolong Glass (Invitrogen) and allowed to cure at room temperature for 48 hours prior to imaging.

### FISH: image acquisition, quantification, and analysis

Samples were imaged on a Nikon AX-R laser scanning confocal microscope with 4-channel GaAsP PMT detector modules. Ovaries were imaged under a 20X objective (N.A.=0.7) on resonant scanning mode using 405, 561, and 640 nm laser channels. We imaged 140 ovarioles from co-infected, mated flies, 172 ovarioles from co-infected, unmated flies, 64 ovarioles from monoinfected, mated flies, and 24 ovarioles from monoinfected, unmated flies. All samples were imaged at Nyquist resolution, with Z-steps calculated accordingly. Image processing and analysis was performed in NIS Elements Advanced software v. 6.10.01. Images were denoised with the FastDenoising tool, deconvoluted with the 3D deconvolution tool, and the backgrounds removed with a rolling ball algorithm (radius=25 pixels). Regions of interest (ROIs) were selected by drawing around individual egg chambers using the polygon tool. Due to high *Wolbachia* density and/or aggregations (see results), individual *Wolbachia* puncta could not be counted in many regions. Instead, we measured the area occupied by each strain inside the egg chamber to infer abundance. Thresholding was used to generate a binary mask representing the area occupied by each strain within each ROI, for each Z-layer. An R script was used to summate the area of the binary layer of each channel in each individual ROI. Egg chambers were staged manually by size and morphology based on Jia, D., Xu, Q., Xie, Q. *et al.* [55]. Stage 1 egg chambers are located within germaria and difficult to differentiate from germarial tissues without additional markers. Thus, germarium measurements here include both the germaria and any stage 1 egg chambers located within them. Embryos were imaged and processed following the same procedures as ovaries, with the only difference being use of a 10X objective (N.A.=0.45). Laser power and acquisition settings were kept consistent across all samples of the same type (i.e., ovary or embryo).

### FISH: statistics and data visualization

Statistical analyses and data visualization were performed in R version 4.4.2 [56]. In all cases we used a generalized linear model (glm) with Gamma variance to model *Wolbachia* abundance. First, the ovaries of unmated and mated females were separately modeled to assess *w*Ha and *w*No abundance with egg chamber stage, strain, and their interaction as fixed effects. To isolate strain-specific impacts due to mating in co-infected ovaries, we used separate glms to model *Wolbachia* abundance (either *w*Ha or *w*No) with egg chamber stage, mating status, and their interaction as fixed effects. To compare the abundance of *w*Ha in egg chambers derived from co-infected versus monoinfected females, we used a glm with egg chamber stage, co-infection, mating status, and their interactions as fixed effects. A small number of zeros were omitted from glms due to use of a Gamma distribution model, but were included on plots. In all cases, post hoc tests were performed with the R package emmeans [57] using estimated marginal means to make pairwise comparisons with Bonferroni’s correction. Curves comparing strain dynamics were generated with stat_smooth using method= “loess”. Data were visualized with ggplot2 [58] and annotated in Inkscape version 1.2 [59].

## RESULTS

### Strain-specific FISH probes facilitate visualization of co-infection dynamics

We developed strain-specific probes targeting 16S. Resequencing revealed that the 16S locus from each strain in our stock was identical to published reference genomes. To test for autofluorescence, co-infected and monoinfected fly ovaries were subjected to the same FISH protocol, but without strain-specific probes. Only very faint, nonspecific autofluorescence was detected from ovary tissue in the probe channels, regardless of *Wolbachia* status of the fly (Supplemental Figures S1, S2). To test for off-target binding of FISH probes, we stained *w*Ha monoinfected ovaries with both probes. Only the *w*Ha-specific probe produced strong signal and visible punctae (Supplemental Figure S3). Additionally, we stained *Wolbachia*-uninfected ovaries and embryos with both probes (Supplemental Figures S4, S5). *Wolbachia*-uninfected, probe-stained ovaries showed faint, nonspecific fluorescence (Supplemental Figure S4). In mid to late embryogenesis, some background autofluorescence was observed in probe-stained, *Wolbachia*-uninfected embryos, primarily in the yolk (Supplemental Figure S5). This autofluorescence could easily be differentiated from true probe signal (seen in the stained, *Wolbachia*-infected embryos, see results below) due to differences in size, shape, location, and intensity. Co-infected ovaries were stained with both *w*Ha- and *w*No-specific probes and confirmed that probe signals do not significantly colocalize (Supplemental Figure S6). Finally, we compared previous whole ovary measurements of *w*Ha and *w*No abundance produced by qPCR [48] with measurements produced by our FISH-based quantification method and found that both methods showed a similarly *w*No-dominated ovary (Supplemental Figure S7) [48].

### Strains *w*Ha and *w*No are distributed differently across oogenesis

We first examined egg chambers from unmated females to observe *w*Ha and *w*No across oogenesis (Figure 1). Beginning with the germarium, *w*Ha was present at much lower abundance than *w*No, which was consistently highly concentrated in the tips of germaria (Figure 2A). Both strains were present in nurse cells and early oocytes. In early stage (1-7) egg chambers, both strains exhibited strong perinuclear localization in nurse cells. Additionally, *w*Ha formed distinct aggregations (Figure 2B). This pattern disappeared by mid-oogenesis (stages 9-10), when both strains assumed a more cytoplasmic distribution in nurse cells and oocytes (Figure 2C). Finally, in late stages (11-14) of oogenesis, when nurse cells transport cytoplasmic contents into the developing oocyte, *w*No was localized to the anterior of the oocyte (Figure 2D). This pattern was consistent, unique to *w*No, and remained throughout the rest of oogenesis, during which *w*Ha exhibited a diffuse cytoplasmic distribution (Figures 1, 2D).

**Figure 1.**
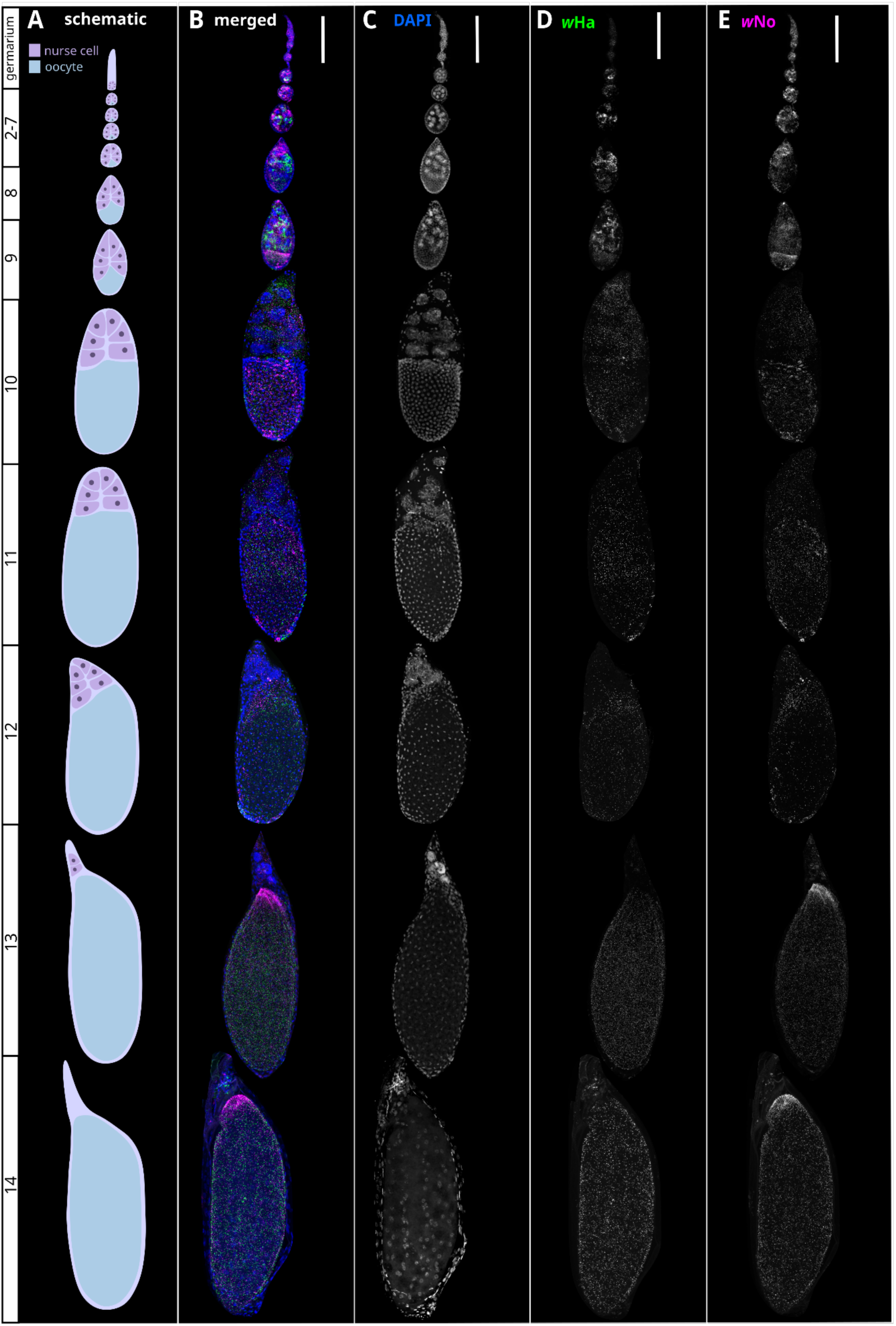
Localization patterns of *w*Ha and *w*No change across oogenesis. Maximum intensity z-projections of egg chambers, stained with strain-specific FISH probes and DAPI. (A) Schematic of a *Drosophila* ovariole showing stages of egg chamber development. (B) Composite representative of a co-infected *Drosophila simulans* ovariole, assembled from images derived from multiple ovarioles, to show *w*Ha and *w*No abundance at each stage of oogenesis. (C) DAPI signal showing egg chamber structure. (D) *w*Ha distribution across oogenesis. (E) *w*No distribution across oogenesis. All scale bars indicate 100 microns. Stage 1 egg chambers are included in germaria.

**Figure 2.**
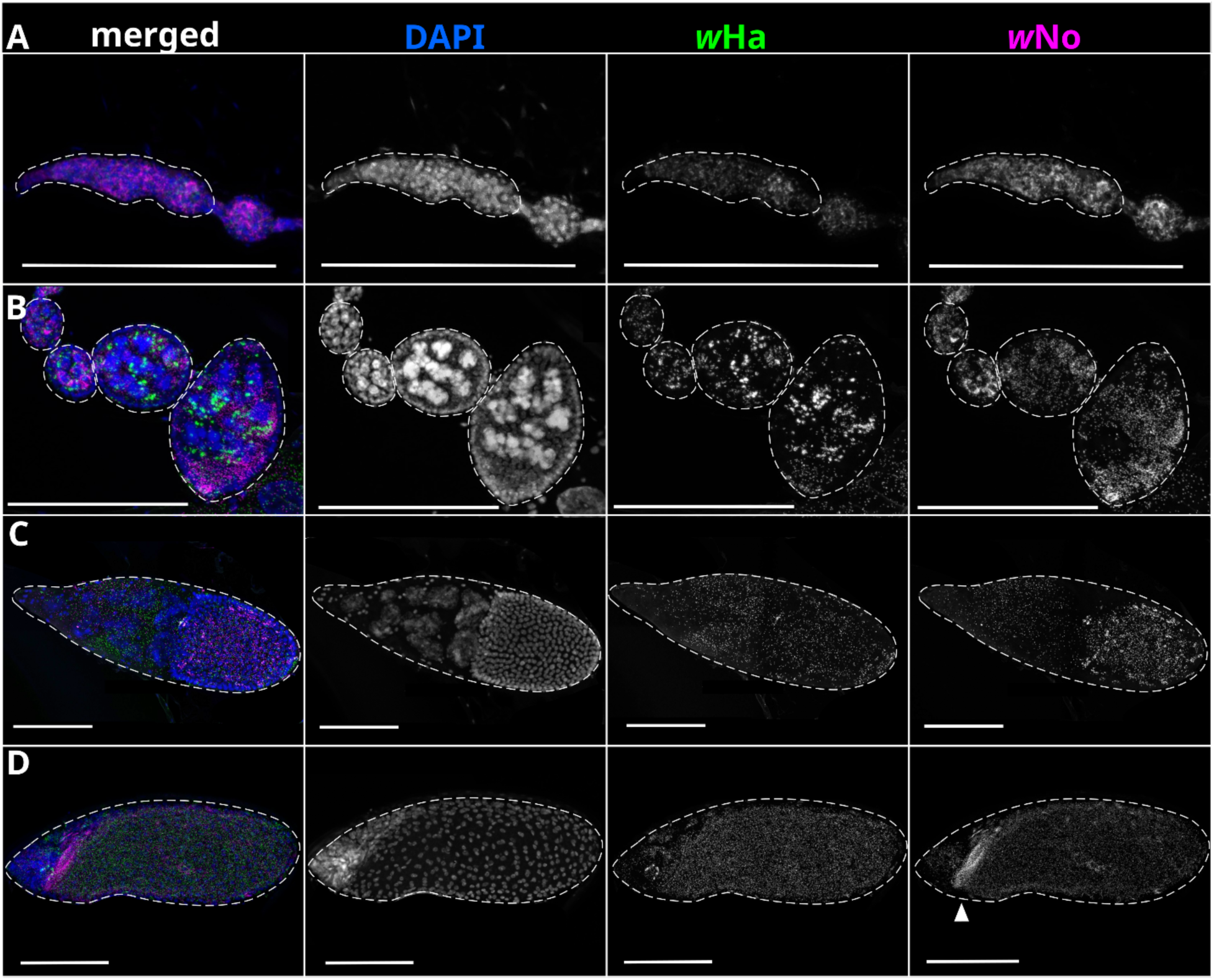
High resolution visualization of co-infection patterns. Maximum intensity z-projections of egg chambers, stained with strain-specific FISH probes and DAPI. Columns from left to right: merged image, greyscale DAPI, greyscale *w*Ha, greyscale *w*No. (A) Germarium and stage 1 egg chamber (dotted outline), with a high concentration of *w*No. (B) Early egg chambers (dotted outlines) with a ‘clumped’ distribution of *w*Ha. (C) Stage 10 egg chamber (dotted outline), with nurse cells and developing oocyte. (D) Stage 12 egg chamber (dotted outline), with nurse cells at the anterior of the egg chamber undergoing apoptosis. Arrowhead indicates the *w*No aggregation at the anterior of the oocyte. All scale bars indicate 100 microns.

### *w*No is the dominant strain throughout oogenesis

To test whether egg chamber development had a significant effect on strain-specific dynamics, we quantified *Wolbachia* abundance from a minimum of 10 replicates of each egg chamber stage, with the exception of stage 11, which was rare (n=1). We determined that the interactive effects of egg chamber stage and strain (β=-1.535e-05± 1.525e-06 SE, t=-10.07, p<0.0001), egg chamber stage (β=-4.686e-06± 3.417e-07 SE, t=-13.71, p<0.0001), and strain (β= 2.299e-04± 2.039e-05 SE, t=11.28, p<0.0001) all significantly impacted *Wolbachia* abundance. Post hoc tests revealed that *w*No was significantly more abundant than *w*Ha in all egg chambers except for in stage 11 (which had low replication), indicating that *w*No remained the more abundant strain in egg chambers throughout oogenesis in ovaries of unmated females.

We wondered if the dominance of *w*No throughout oogenesis (Figure 3A) was specific to unmated females, and if the co-infection in late-stage egg chambers from mated females would align more with the patterns we saw in early stages of embryogenesis wherein *w*Ha and *w*No had more similar genome copy numbers [48]. To address this, we examined *w*Ha and *w*No abundance in the ovaries of mated females (Figure 3B) and determined that the interactive effects of egg chamber stage and strain (β=-1.108e-05± 1.274e-06 SE, t=-8.695, p<0.0001), egg chamber stage (β=-3.859e-06± 2.976e-07 SE, t=-12.964, p<0.0001), and strain (β= 1.714e-04± 1.662e-05 SE, t=10.314, p<0.0001) all significantly impacted *Wolbachia* abundance. Post hoc tests revealed that *w*No was significantly more abundant than *w*Ha in all egg chambers except for those in stage 6 and 11 (which had low replicates, n=5 and n=2, respectively), indicating that *w*No was also the more abundant strain in ovaries of mated females.

**Figure 3.**
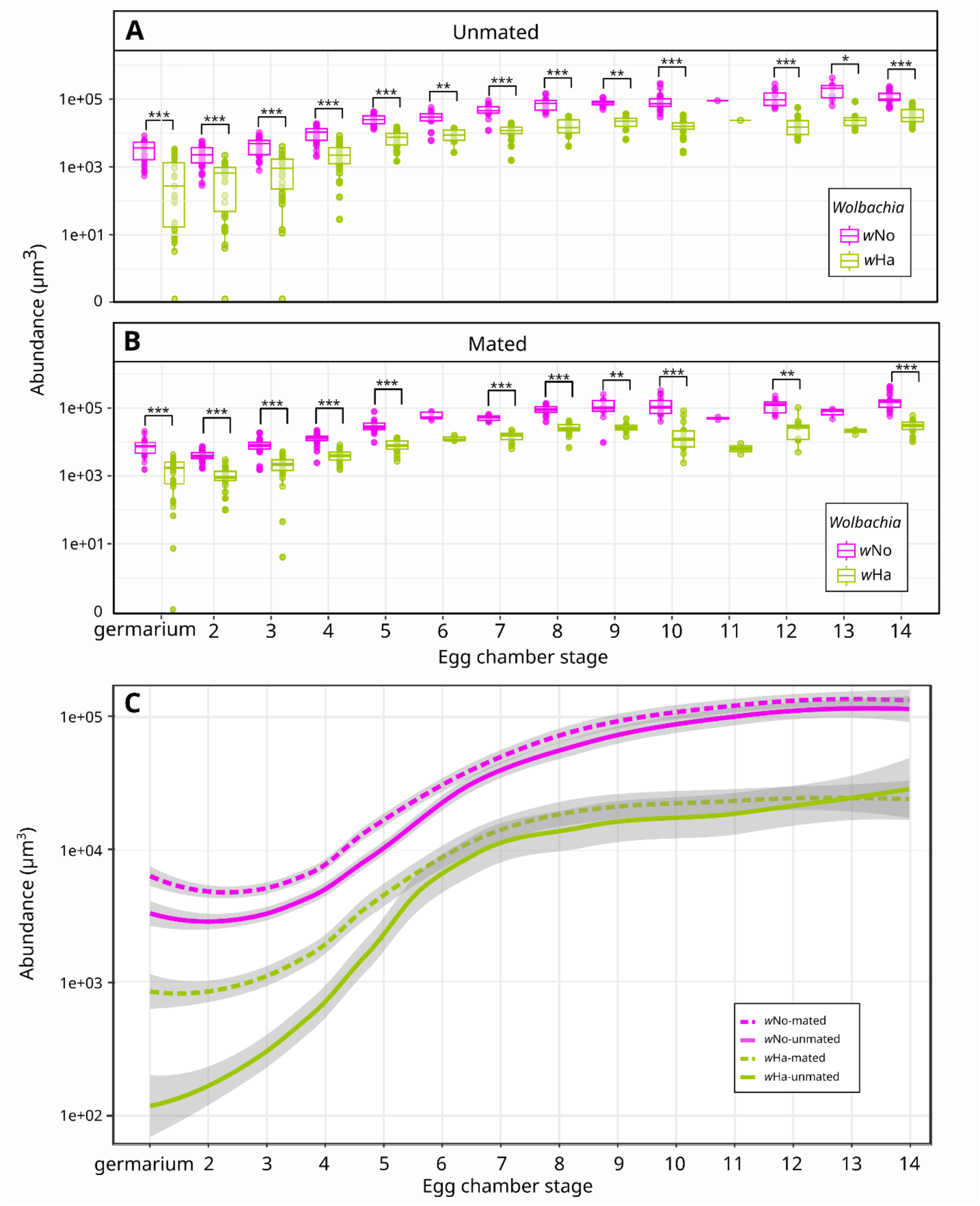
Strain-specific *Wolbachia* abundance changes across development, is sensitive to mating, and maintains a *w*No bias. Abundance was calculated from summed area of fluorescent signal detected for each probe channel in individual egg chambers from ovaries stained with strain-specific probes. (A) Abundance of *w*Ha and *w*No in ovaries of unmated females. (B) Abundance of *w*Ha and *w*No in ovaries of mated females. (C) Trend lines showing the change in abundance of each strain during oogenesis for mated and unmated females. Standard error was calculated for a 95% confidence interval. Significance values ***<0.001, **<0.01, *<0.05 calculated from estimated marginal means test with Bonferroni correction for multiple comparisons. Stage 1 egg chambers are included in germaria. A small number of zeros ((A) n=5 *w*Ha, (B) n=1 *w*Ha) were omitted from statistical tests, but included in plots.

### Abundance of both strains in early egg chambers increases upon mating

While *w*No was more abundant across oogenesis regardless of whether females were mated, because *Drosophila* ovary maturation is completed in response to mating [60–62] we wondered if mating influenced strain-specific dynamics. First, we examined *w*No abundance. The interaction of egg chamber stage and mating status was not significant (β=-8.277e-07 ± 4.536e-07 SE, t=-1.825, p=0.0686), but egg chamber stage (β=-3.859e-06 ± 2.907e-07 SE, t=-13.273, p< 0.0001) and mating status (β=1.219e-05 ± 6.070e-06 SE, t=2.008, p=0.0452) alone had significant effects on *w*No abundance. Post-hoc tests revealed that *w*No abundance was significantly higher in germaria (adj. p=0.0309), stage 2 (adj. p=0.0003), and stage 3 (adj. p<0.0001) egg chambers from mated females, indicating that mating significantly increased *w*No abundance in early oogenesis (Figure 3C).

Next, we examined the effects of mating, egg chamber stage, and their interaction on *w*Ha abundance. Similarly, we found that the interaction of egg chamber stage and mating status (β=-5.101e-06 ± 1.948e-06 SE, t=-2.618, p=0.0091), egg chamber stage (β=-1.494e-05 ± 1.216e-06 SE, t=-12.282, p<0.0001), and mating status (β=7.065e-05 ± 2.578e-05 SE, t=2.740, p=0.0063) all had significant effects on *w*Ha abundance. Post-hoc tests revealed that *w*Ha was significantly more abundant in germaria (adj. p=0.011), stage 2 (adj. p=0.0031), and stage 3 egg chambers from mated females (adj. p=0.0044) indicating that mating also increased *w*Ha abundance in early egg chambers (Figure 3C).

### *w*Ha and *w*No localize differently across embryonic development

Given the different distributions of *w*Ha and *w*No we observed in mature oocytes, we next assessed how *w*Ha and *w*No were distributed in embryos (Figure 4). Very early in development, prior to cellularization, *w*No remained strongly localized to the anterior pole, but was also weakly aggregated around nuclei and at the peripheral surface of the embryo (Figure 3A). At this time, *w*Ha had a strong perinuclear localization. Immediately following cellularization, *w*No remained at a higher density at the anterior pole but was also present at high titers throughout the embryo (Figure 4B). Neither *w*Ha nor *w*No maintained the perinuclear localization patterns observed prior to cellularization. Both strains were present in pole cells and there was a very subtle enrichment of both strains in that region (Figure 4B). Next, we examined embryos during the stages at which pole cells migrate. The anterior concentration of *w*No observed in previous stages dissipated by stage 10 of embryogenesis (Figure 4C). Aggregations of *w*Ha and *w*No were clearly present in the region where pole cells are expected to reside at this stage (Figure 4C). Finally, we examined embryos after formation of the gonadal imaginal discs. *w*Ha and *w*No were concentrated in bilaterally symmetrical aggregations located where we would expect gonadal imaginal discs to reside (Figure 4D).

**Figure 4.**
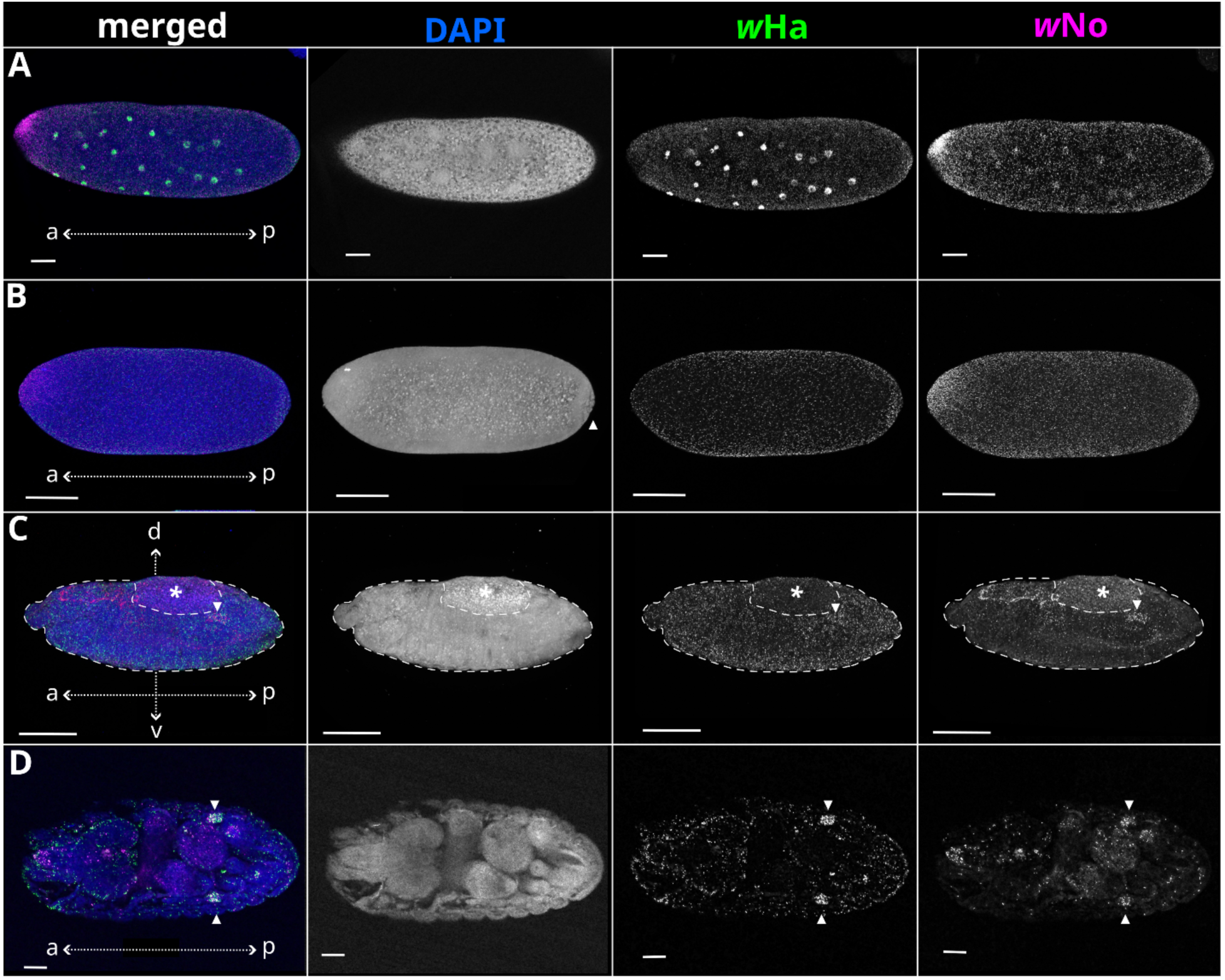
Distribution of co-infecting *Wolbachia* strains changes across embryo development. Maximum intensity z-projections of embryos, stained with strain-specific FISH probes and DAPI. Columns from left to right: merged image, greyscale DAPI, greyscale *w*Ha, greyscale *w*No. Dotted arrows indicate embryo anterior-posterior and dorsal-ventral axes (a, anterior; p, posterior; d, dorsal; v, ventral). (A) Stage 3 embryo, with anterior localization of *w*No and perinuclear localization of *w*Ha (B) Stage 5 embryo, undergoing formation of cellular blastoderm and pole cells. Arrowhead indicates location of pole cells. (C) Stage 14 embryo (dotted outline), undergoing dorsal closure with *Wolbachia* localized to the putative gonadal imaginal discs (arrowhead). Asterisk indicates location of autofluorescent yolk. (D) Stage 16 embryo, with localization of *w*Ha and *w*No to the putative gonadal imaginal discs (arrowheads). All scale bars indicate 100 microns.

### Transmission of *w*No fails in the absence of *w*Ha

We successfully established stable *w*Ha monoinfected isofemale lines (n=3), all of which stably transmitted *w*Ha for at least five generations. A single *w*Ha monoinfected line was selected for continued maintenance and has since been kept in culture for >130 generations and still stably transmits *w*Ha with 100% fidelity. In contrast, *w*No segregants were extremely rare (<1% of screened flies) and within two generations all isofemale lines initiated from *w*No monoinfected females (n=3) produced only *Wolbachia*-uninfected offspring. Due to this transmission failure, we were unable to generate a stable *w*No monoinfected line. Furthermore, isofemale lines with co-infections that were derived from heat-treated mothers (n=3) all stably transmitted both *w*Ha and *w*No for at least 3 generations.

### *w*Ha infection dynamics depend on the presence of *w*No and mating

Clearly, *w*No is severely impacted by the absence of *w*Ha. While *w*Ha transmission is not destabilized by the absence of *w*No, we wondered whether *w*Ha abundance and localization patterns change during monoinfection. Thus, we examined egg chambers and embryos of *w*Ha moninfected flies and compared these patterns with those of *w*Ha in the above-described co-infected flies. In the absence of *w*No, *w*Ha maintains a clustered distribution in early egg chambers and diffuse localization pattern in late-stage egg chambers (Figure 5). We also identified no clear differences in the distribution of *w*Ha in monoinfected embryos (Supplemental Figure S8). While *w*Ha tropism appears to be independent of co-infection with *w*No, we considered that the abundance of *w*Ha could differ in the absence of *w*No. We compared the abundance of *w*Ha in egg chambers derived from co-infected versus monoinfected females and found that the interaction of egg chamber stage, mating status, and co-infection (β= -1.370e-05± 2.707e-06 SE, t=-5.060, p<0.0001), the interaction of mating status and co-infection (β= 1.987e-04± 3.526e-05 SE, t= 5.635, p<0.0001), the interaction of egg chamber stage and co-infection (β= 6.035e-06± 1.871e-06 SE, t= 3.226, p=0.0013), the interaction of egg chamber stage and mating status (β= 8.596e-06± 2.087e-06 SE, t= 4.118, p<0.0001), egg chamber stage (β= - 2.097e-05± 1.531e-06 SE, t= -13.703, p<0.0001), mating status (β= -1.281e-04± 2.689e-05 SE, t= -4.762, p<0.0001), and co-infection (β= -8.762e-05± 2.419e-05 SE, t= -3.622, p=0.0003) all had significant effects on *w*Ha abundance. Specifically, in unmated females, *w*Ha was more abundant in monoinfected germaria (adj. p<0.0001), stage 2 (adj.p<0.0001), and stage 4 egg chambers (adj. p=0.0111). In contrast, in mated females, *w*Ha was more abundant in monoinfected germaria (adj. p=0.0141) but less abundant in monoinfected stage 8 egg chambers (adj. p=0.0069). These findings suggest that *w*Ha abundance differs across oogenesis depending on both co-infection and the mating status of the female.

**Figure 5.**
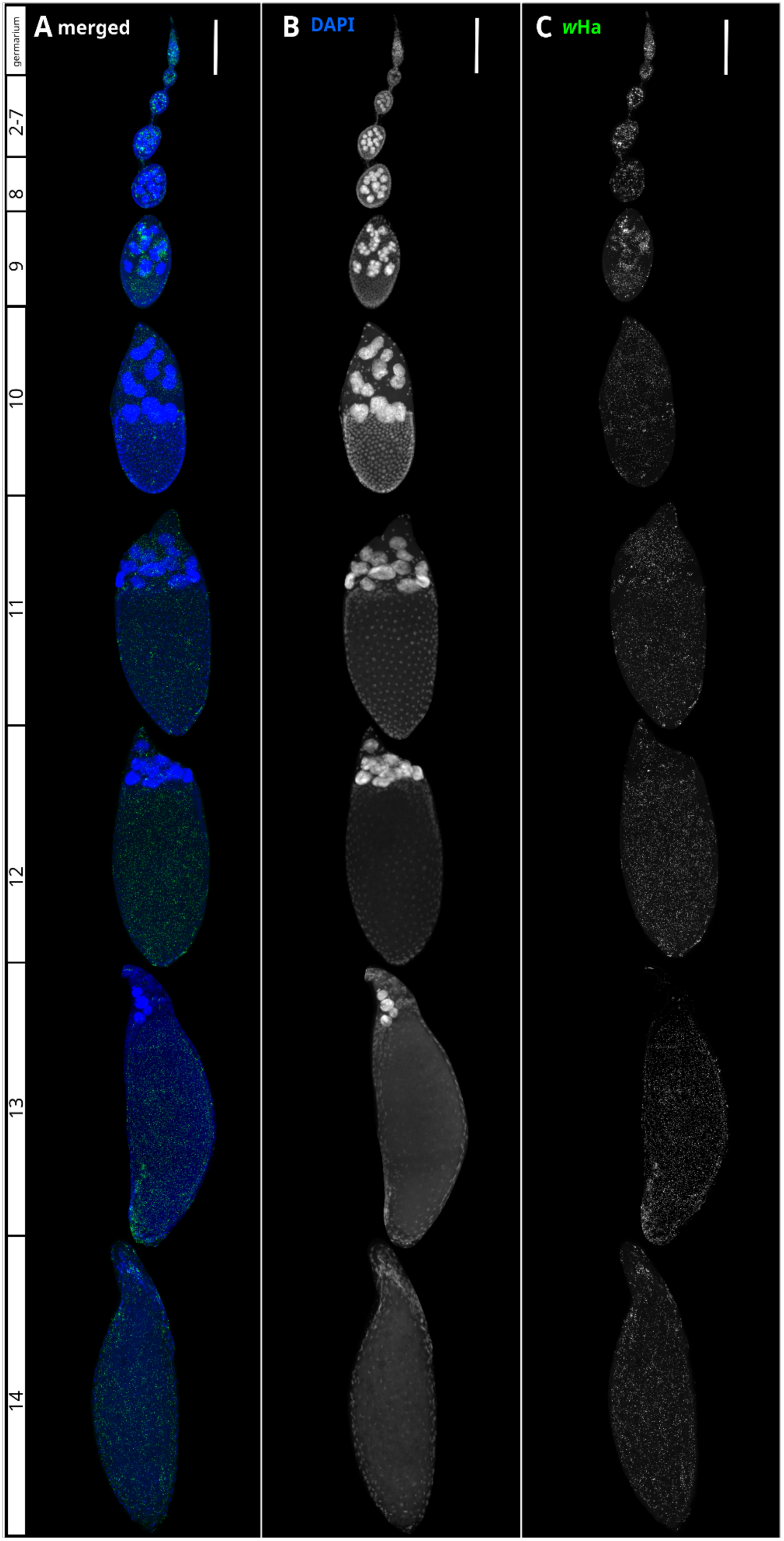
*w*Ha distribution patterns in ovarioles in the absence of *w*No. Maximum intensity z-projections of egg chambers, stained with strain-specific FISH probes and DAPI. (A) Composite representative of a *w*Ha monoinfected *Drosophila simulans* ovariole, assembled from multiple images to show stages of egg chamber development. Egg chamber stages are shown on the left. (B) Greyscale DAPI signal showing egg chamber structure. (C) Greyscale *w*Ha distribution. All scale bars indicate 100 microns. Stage 1 egg chambers are included in germaria.

**Figure 6.**
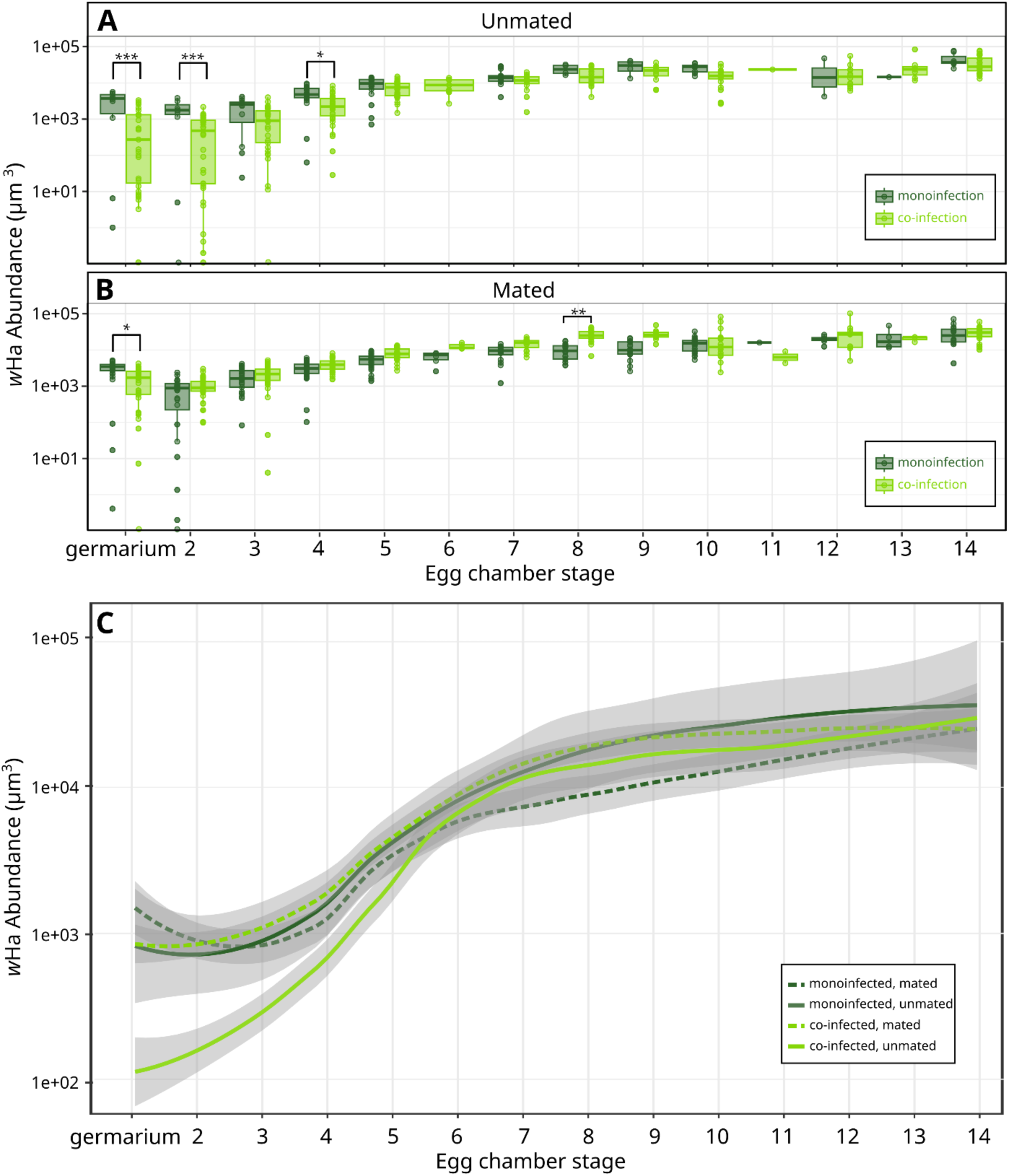
*w*Ha abundance dynamics differ between co-infected and monoinfected ovaries. (A) Abundance of *w*Ha in ovaries of unmated females throughout oogenesis. (B) Abundance of *w*Ha in ovaries of mated females across oogenesis. (C) Curves comparing dynamics of *w*Ha showing effects of mating and co-infection. Standard error calculated for a 95% confidence interval. Significance values ***<0.001, **<0.01, *<0.05 calculated from estimated marginal means test with Bonferroni correction for multiple comparisons. Stage 1 egg chambers are included in germaria. A small number of zeros ((A) n=1 monoinfected, n=5 co-infected, (B) n=1 monoinfected, n=1 co-infected) were omitted from statistical tests, but included in plots.

## DISCUSSION

Understanding how multiple intracellular symbionts achieve vertical transmission through the germline, while competing for limited space and resources has important implications for studying the ecology, evolution, and life history of intracellular organisms [63]. Additionally, improved understanding of stabilizing factors present in natural co-infections will aid in the use of *Wolbachia*-based methods to manage populations of agricultural pests and vector species, particularly those that are naturally host to one or more *Wolbachia* strains [64, 65]. Lessons learned from stably transmitted natural co-infections may help guide targeted artificial establishment of co-infections for *Wolbachia*-based population replacement programs, which are reliant on stable transmission to be effective [35, 65, 66].

Other *Wolbachia* exhibit strain-specific differences in ovary tissue tropism [42] and spatial dynamics in oocytes [44] and embryos [46]. We suggest that co-infection stability might be facilitated by avoiding inter-strain competition through occupying distinct niches in the host. Our results indicate that co-infecting *Wolbachia* strains *w*Ha and *w*No do indeed have some biases for different spatial niches across oogenesis. A striking example of this is the aggregation of *w*No at the anterior pole during a stage that coincides with nurse cell apoptosis and transport of nurse cell contents, including *Wolbachia,* into the oocyte [67, 68]. Despite clear spatial differences between strains appearing in late oogenesis, *w*No remained significantly more abundant than *w*Ha throughout the entirety of oogenesis.

We hypothesized that strain-specific differences in spatial occupancy of ovary tissues may reconcile the apparent discrepancy in relative abundances of *Wolbachia* strains between ovaries and two hour old embryos we observed previously [48]. For example, if perhaps *w*No preferentially occupied early-stage egg chambers, but only weakly targeted the mature oocyte, *w*No could be the more abundant strain in whole ovaries, whilst being transmitted at a much lower relative abundance and thus less abundant in embryos. However, this does not appear to be the case, as *w*No is present within developing egg chambers and oocytes at significantly higher titers than *w*Ha throughout the entirety of oogenesis, and *w*No is highly abundant in embryos up to cellularization. This discrepancy might reflect differences in the biology of each strain that impact quantification. For example, genome endoreplication would affect qPCR-based quantification, while differences in cell size or concentration of 16S rRNA transcripts would affect our FISH-based quantification method. Regardless, there are still clear strain-specific shifts in infection dynamics during embryo development that indicate *w*Ha and *w*No have very different approaches to interacting with host biology. For example, similar to the late-stage oocytes, a significant population of *w*No remains concentrated at the anterior pole of the embryo until around cellularization. Conversely, *w*Ha strongly localizes perinuclearly during early syncytial divisions. It is unclear what host structures or proteins, each strain is associating with, though the distribution of *w*No resembles that of *bicoid* and some other maternally provisioned axis patterning transcripts that are anchored at the anterior oocyte cortex [68, 69]. As embryogenesis progresses, spatial partitioning between *w*Ha and *w*No becomes less apparent. Following dorsal closure, both strains are concentrated in what are likely the developing gonadal imaginal discs, based on their size and location [70], though further investigation with a germline-specific marker will be valuable for testing this hypothesis.

In addition to navigating spatio-temporal aspects of host development, these *Wolbachia* strains must navigate mating-induced structural and metabolic changes that occur in *Drosophila* ovaries. Upon mating, *Drosophila* germline stem cells proliferate [61, 62] and vitellogenesis resumes to accommodate an increased rate of oogenesis [60]. *Wolbachia* may compensate for the increased turnover in the ovaries by increasing replication rate in the germline or invading from nearby somatic tissue. We show that strains are differently impacted by mating-associated changes in the ovaries and thus may employ different strategies to maintain an adequate pool of *Wolbachia* for transmission. Interestingly, not only was *w*Ha impacted by mating, but the combination of an unmated host and the presence of *w*No led to significantly lower *w*Ha abundance in early stages of oogenesis. The biological relevance of these co-infection sensitive responses to mating are unclear. *w*Ha may simply increase in abundance to capitalize on the intracellular space or nutrients made available by the absence of *w*No. Alternatively, mating and co-infection sensitive effects on *w*Ha abundance may be a result of more complex direct interactions with *w*No, the host, or both.

While we found that *w*Ha is transmitted reliably in flies cleared of *w*No, *w*No transmission is severely impacted by the loss of *w*Ha, despite *w*No’s greater abundance. Alternatively, *w*No may be more sensitive to heat than *w*Ha, and transgenerational effects of heat shock manifest as transmission failure. We consider this hypothesis unlikely, considering co-infected isofemale lines derived from heat-shocked mothers reliably transmit *w*No over many generations. If differential sensitivity to heat treatment is indeed responsible for transmission failure of *w*No, it appears that the presence of *w*Ha mitigates this effect, again supporting the idea of an asymmetric interaction. Given the anterior localization patterns of *w*No in mature oocytes and early embryos, we suggest that transmission failure may be related to an inability to target the posterior pole plasm in sufficient quantities in the absence of *w*Ha. Anterior localization in the oocyte has been observed in other *Wolbachia* strains that are closely related to *w*No [71], though these strains are stably transmitted as monoinfections, indicating that concentration of *Wolbachia* at the anterior pole is not the sole cause of transmission failure in *w*No. Critically, it remains unknown how *w*No localization patterns are altered in the absence of *w*Ha and what the mechanism for rescue by *w*Ha may be. Perhaps *w*No has lost genes for important effector proteins and is reliant on *w*Ha to make and secrete effectors necessary for stable transmission [72]. Importantly, *w*No monoinfections have been isolated in *Drosophila simulans* previously [33, 47, 73], but fly genotypes, strain segregation methods, rearing conditions, and experimental shifts in the intensity of cytoplasmic incompatibility pressure all varied which precludes direct comparisons. Alternative approaches to segregating the two strains, rapid identification of *w*No monoinfected females for examination of their *w*No distributions, as well as sequencing of the *w*No genome in our specific co-infected *Drosophila simulans* could shed light on potential mechanisms behind the transmission failure and differences between experimental results.

In conclusion, we characterized the transmission dynamics of two stably transmitted, co-infecting *Wolbachia* strains in *Drosophila simulans*, identified patterns of niche partitioning, mapped the abundance of both strains across the entirety of oogenesis, and identified strain-specific responses to mating and co-infection. Although we determined that some level of niche partitioning is likely at play, it is unclear why different *Wolbachia* strains have different distributions within egg chambers and embryos. What drives their differences in tropism? What host structures or intracellular areas do they associate with? Do they have different metabolic needs? Future exploration of co-infection dynamics in other models, such as *Nasonia vitripennis* (*w*VitA, *w*VitB) and *Aedes albopictus* (*w*AlbA, *w*AlbB), will enable us to investigate if there are any conserved features associated with stably transmitted *Wolbachia* co-infections. Our work demonstrates that co-transmitted *Wolbachia* pose interesting questions about ecological interactions at the micro scale and represent a novel system to investigate these questions.

## AUTHOR CONTRIBUTIONS

M.W.J. helped design the study, performed experiments and analyses, and wrote the paper. P.A.S. performed experiments. A.R.I.L. designed the study, provided guidance on experimental setup and analysis methods, and assisted in paper writing. All authors contributed to the research and approved the final manuscript.

## ACKNOWLEDGEMENTS

Imaging and analysis were supported by resources and staff at the University of Minnesota University Imaging Centers (UIC) SCR_020997.

## SUPPLEMENTAL FIGURES

**Supplemental Figure S1.**
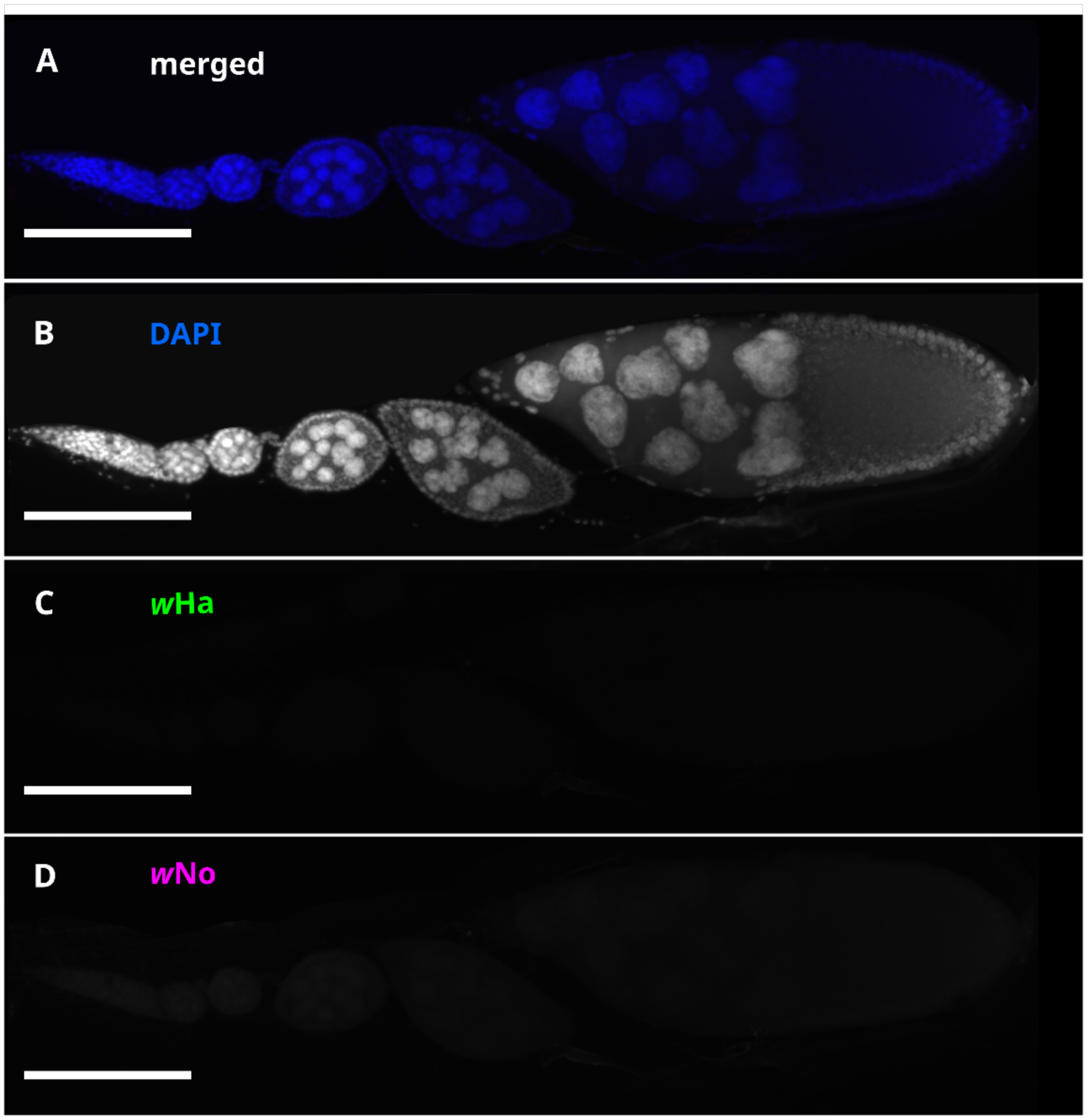
Co-infected, no-probe ovary controls have no fluorescence in the probe channels. Maximum intensity z-projections of egg chambers, stained with DAPI. Control ovaries went through the same FISH sample preparation protocol, with the only difference being the omission of strain-specific probes during the hybridization step. These images had their backgrounds removed with a rolling ball algorithm, but were not denoised or deconvoluted because these processing steps introduce artifacts in channels with limited signal. (A) Merged channels showing no punctae associated with probe channels. (B) Greyscale DAPI signal showing egg chamber structure. (C) Greyscale *w*Ha channel showing a lack of autofluorescent signal and no visible punctae. (D) Greyscale *w*No channel showing lack of autofluorescent signal and no visible punctae. All scale bars indicate 100 microns.

**Supplemental Figure S2.**
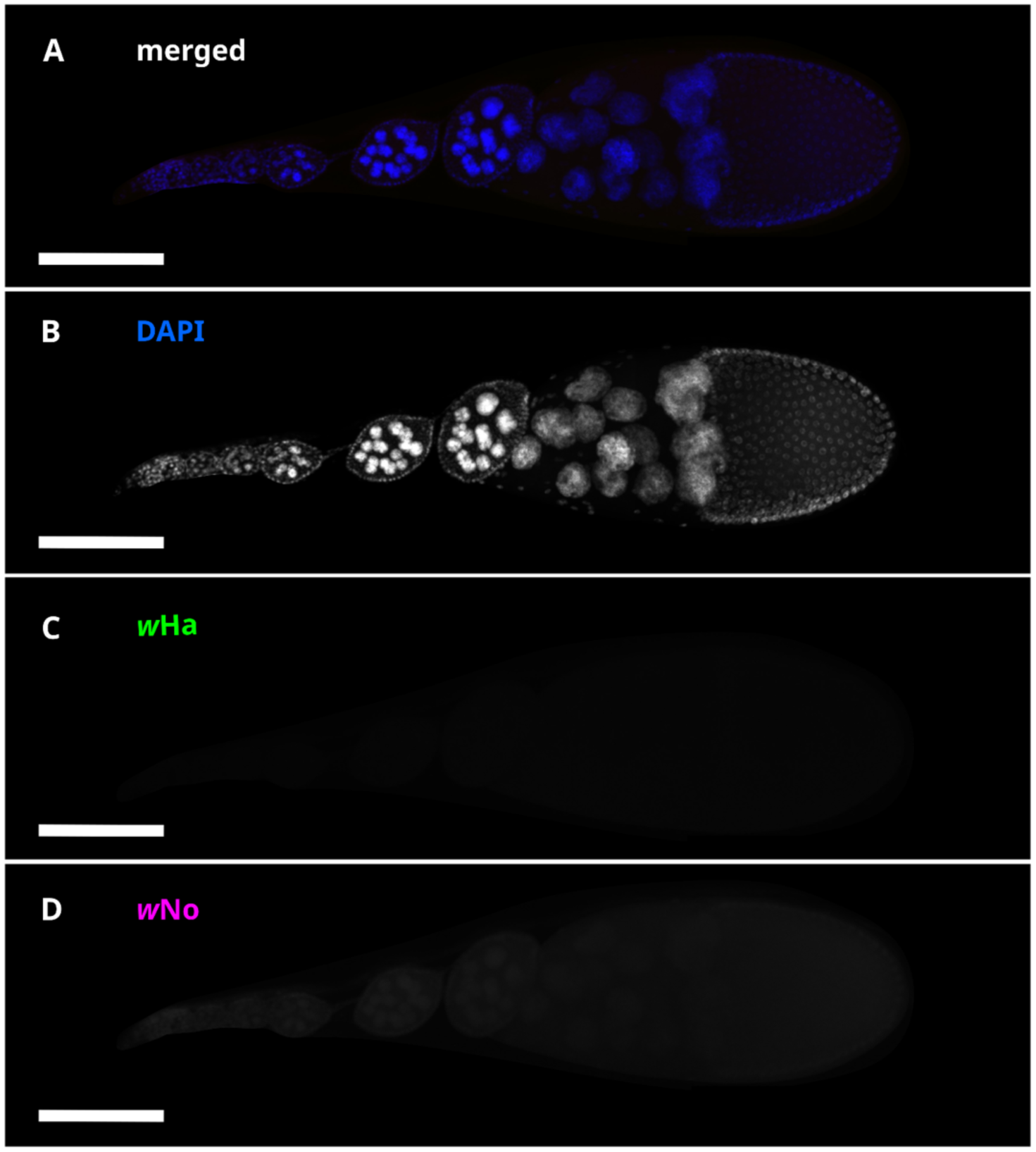
***w*Ha moninfected, no-probe ovary controls have no fluorescence in the probe channel.** Maximum intensity z-projections of egg chambers, stained with DAPI. Control ovaries went through the same FISH sample preparation protocol as ovaries used in experiments, with the only difference being the omission of strain-specific probes during the hybridization step. These images had their backgrounds removed with a rolling ball algorithm, but were not denoised or deconvoluted because these processing steps introduce artifacts in channels with limited signal. (A) Merged channels showing no punctae associated with probe channels. (B) Greyscale DAPI signal showing egg chamber structure, (C) Greyscale *w*Ha channel showing a lack of autofluorescent signal and no visible punctae. (D) Greyscale *w*No channel showing lack of autofluorescent signal and no visible punctae. All scale bars indicate 100 microns.

**Supplemental Figure S3.**
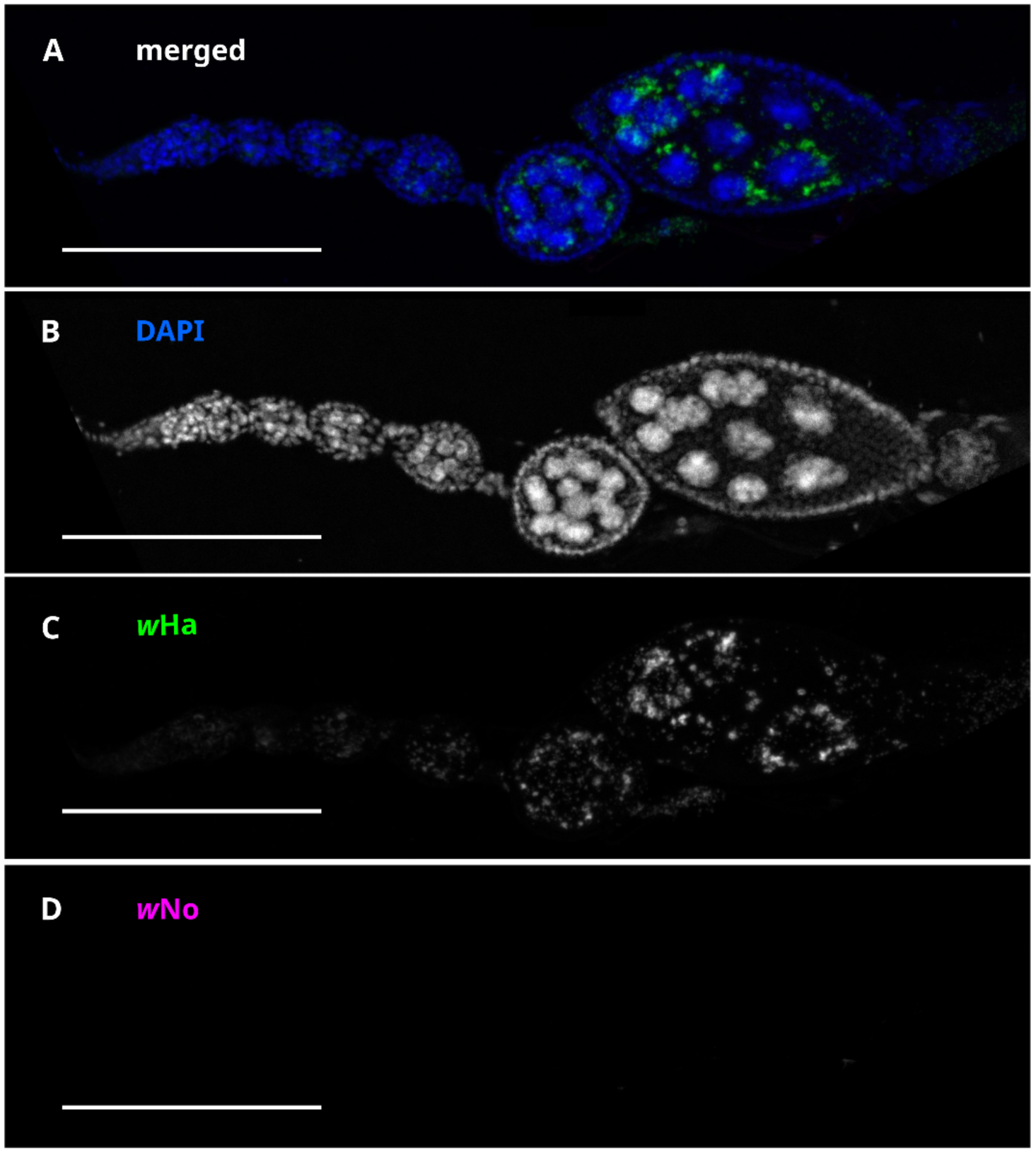
***w*Ha monoinfected ovariole has punctae in the *w*Ha channel and no fluorescence in the *w*No channel.** Maximum intensity z-projections of egg chambers, stained with strain-specific FISH probes and DAPI. Control ovaries went through the same FISH sample preparation and staining protocol as ovaries used in experiments. These images had their backgrounds removed with a rolling ball algorithm, but were not denoised or deconvoluted because these processing steps introduce artifacts in channels with limited signal. (A) Merged channels showing specific staining of *w*Ha by *w*Ha-specific probe and absence of nonspecific staining by *w*No-specific probe. (B) Greyscale DAPI signal showing egg chamber structure. (C) Greyscale *w*Ha channel showing visible punctae. (D) Greyscale *w*No channel showing lack of autofluorescent signal and no visible punctae. All scale bars indicate 100 microns.

**Supplemental Figure S4.**
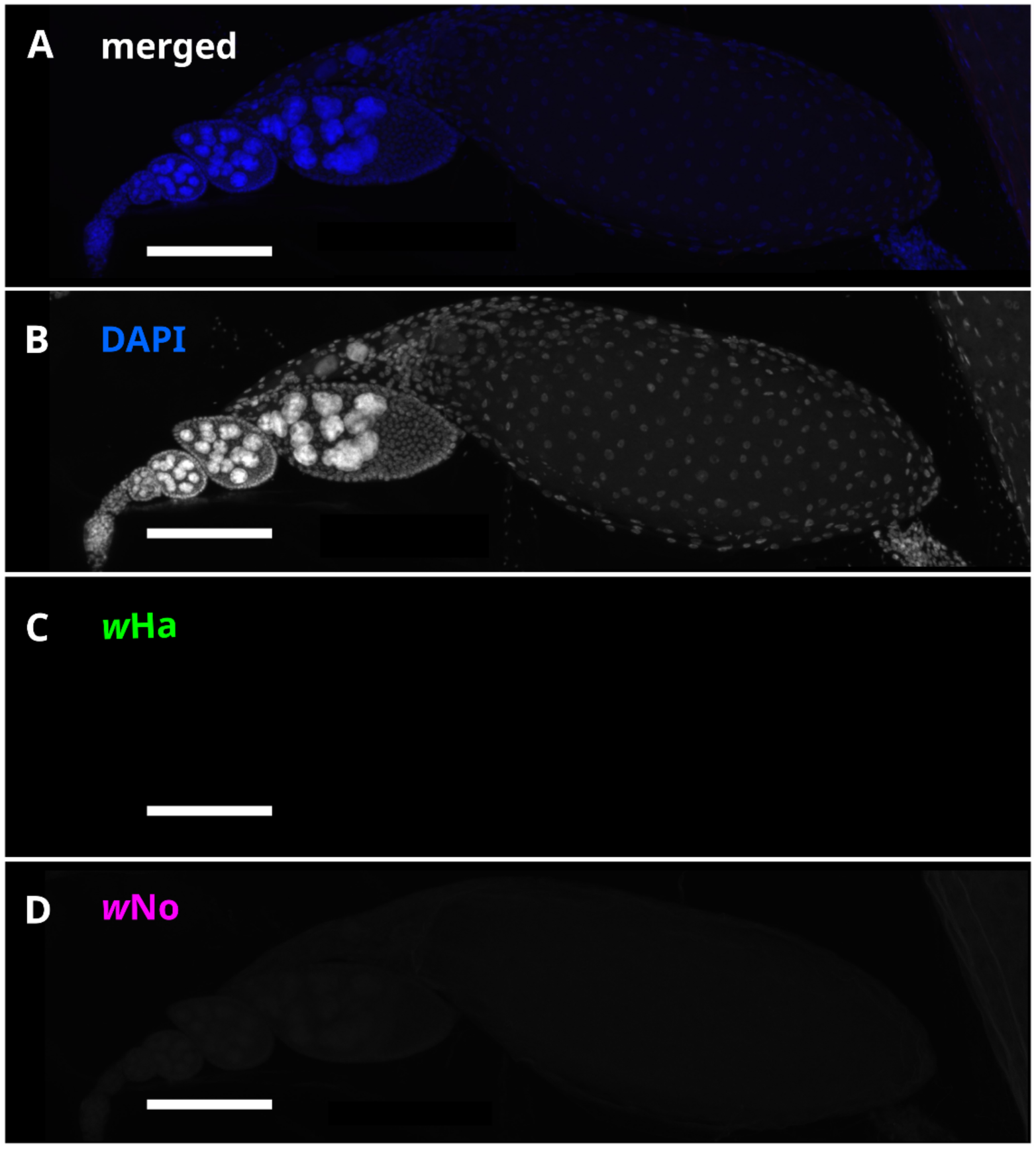
***Wolbachia*-uninfected ovary controls have no fluorescence in either probe channel.** Maximum intensity z-projections of egg chambers, stained with strain-specific FISH probes and DAPI. Control ovaries went through the same FISH sample preparation and staining protocol as ovaries used in experiments. These images had their backgrounds removed with a rolling ball algorithm, but were not denoised or deconvoluted because these processing steps introduce artifacts in channels with limited signal. (A) Merged images showing no nonspecific staining by either probe. (B) Greyscale DAPI signal showing egg chamber structure. (C) Greyscale *w*Ha channel showing no visible punctae. (D) Greyscale *w*No channel showing no visible punctae. All scale bars indicate 100 microns.

**Supplemental Figure S5.**
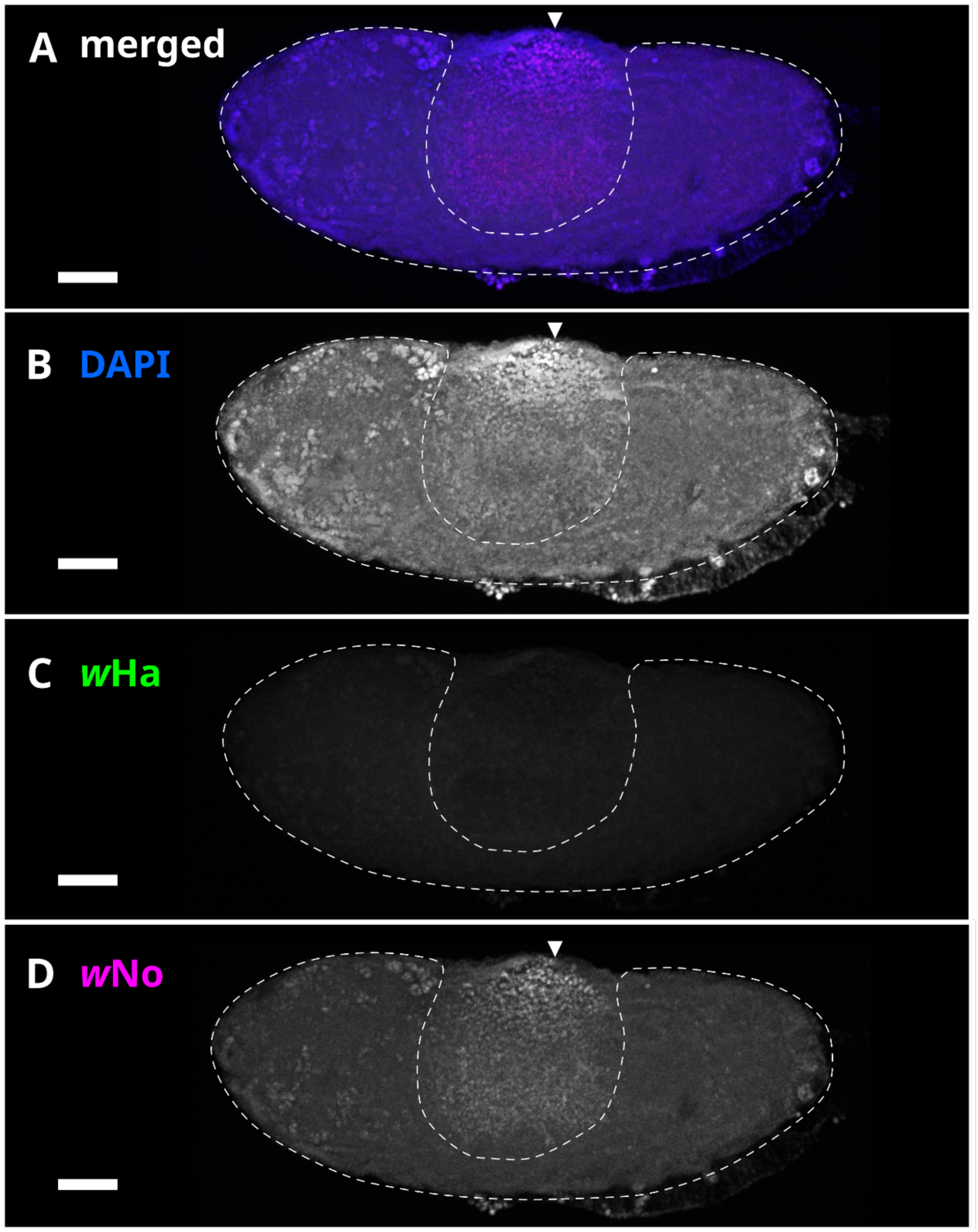
***Wolbachia*-uninfected embryo FISH control.** Maximum intensity z-projections of an embryo, stained with strain-specific FISH probes and DAPI. Control embryos went through the same FISH sample preparation and staining protocol as embryos used in experiments. These images had their backgrounds removed with a rolling ball algorithm, but were not denoised or deconvoluted because these processing steps introduce artifacts in channels with limited signal. The developing embryo is indicated with a dotted outline, and the yolk is indicated with an arrowhead. (A) Merged image of all channels. (B) Greyscale DAPI signal. (C) Greyscale *w*Ha channel. (D) Greyscale *w*No channel with diffuse fluorescence. Yolk granules can be differentiated from true probe signal by comparison of size, shape, location, and intensity of fluorescent signal between *Wolbachia*-uninfected and *Wolbachia*-infected embryos stained with both probes. All scale bars indicate 100 microns.

**Supplemental Figure S6.**
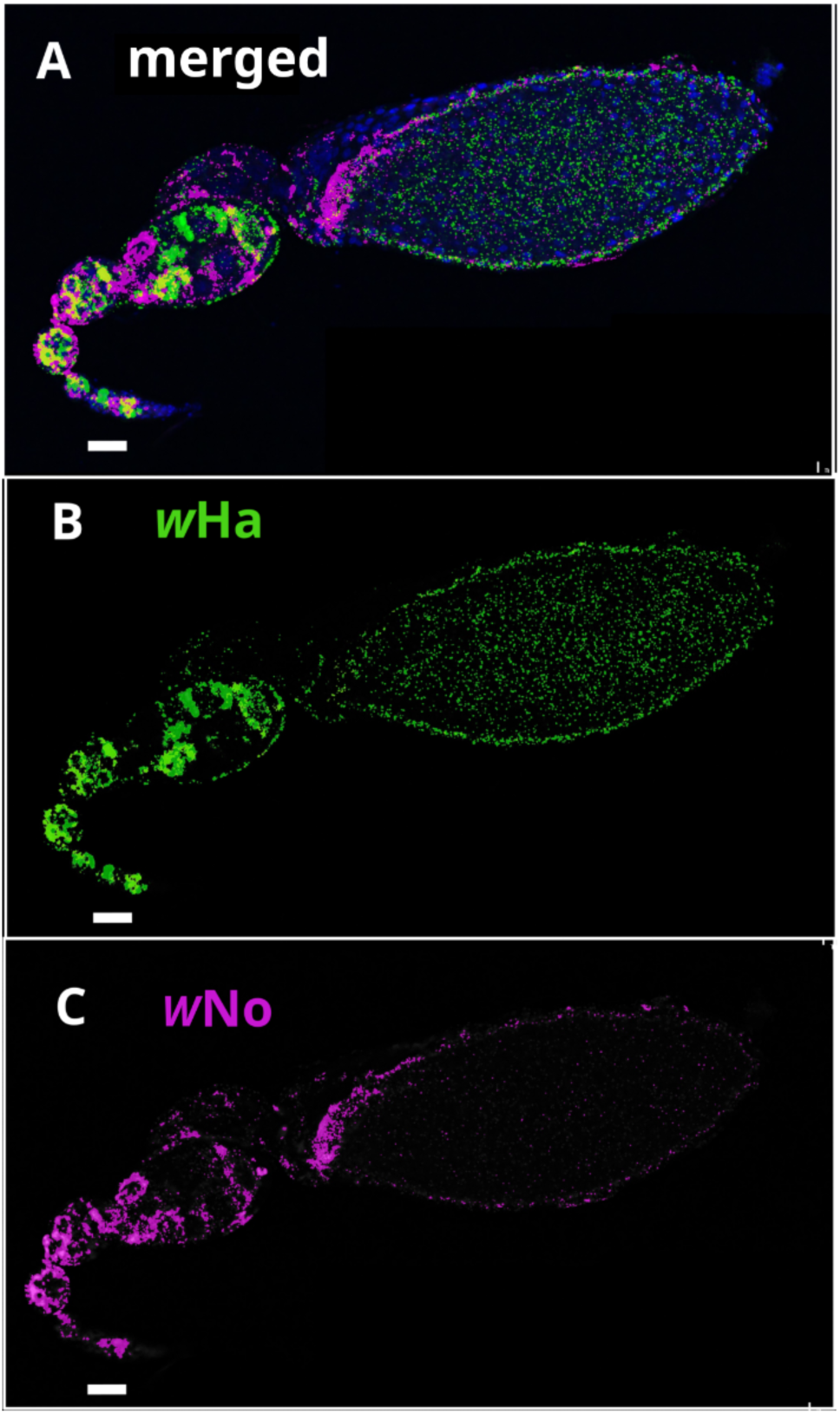
***w*Ha and *w*No probes are strain-specific and signals do not significantly colocalize with one another.** Single layer in z-stack image selected to demonstrate colocalization measurements. (A) Binary mask applied to *w*Ha channel, note that this mask is also used for quantification. (B) Binary mask applied to *w*No channel, note that this mask is also used for quantification. (C) Merged image showing binary masks applied to *w*Ha and *w*No channels, showing overlap between signals in yellow. (Mander’s overlap coefficient=0.44).

**Supplemental Figure S7.**
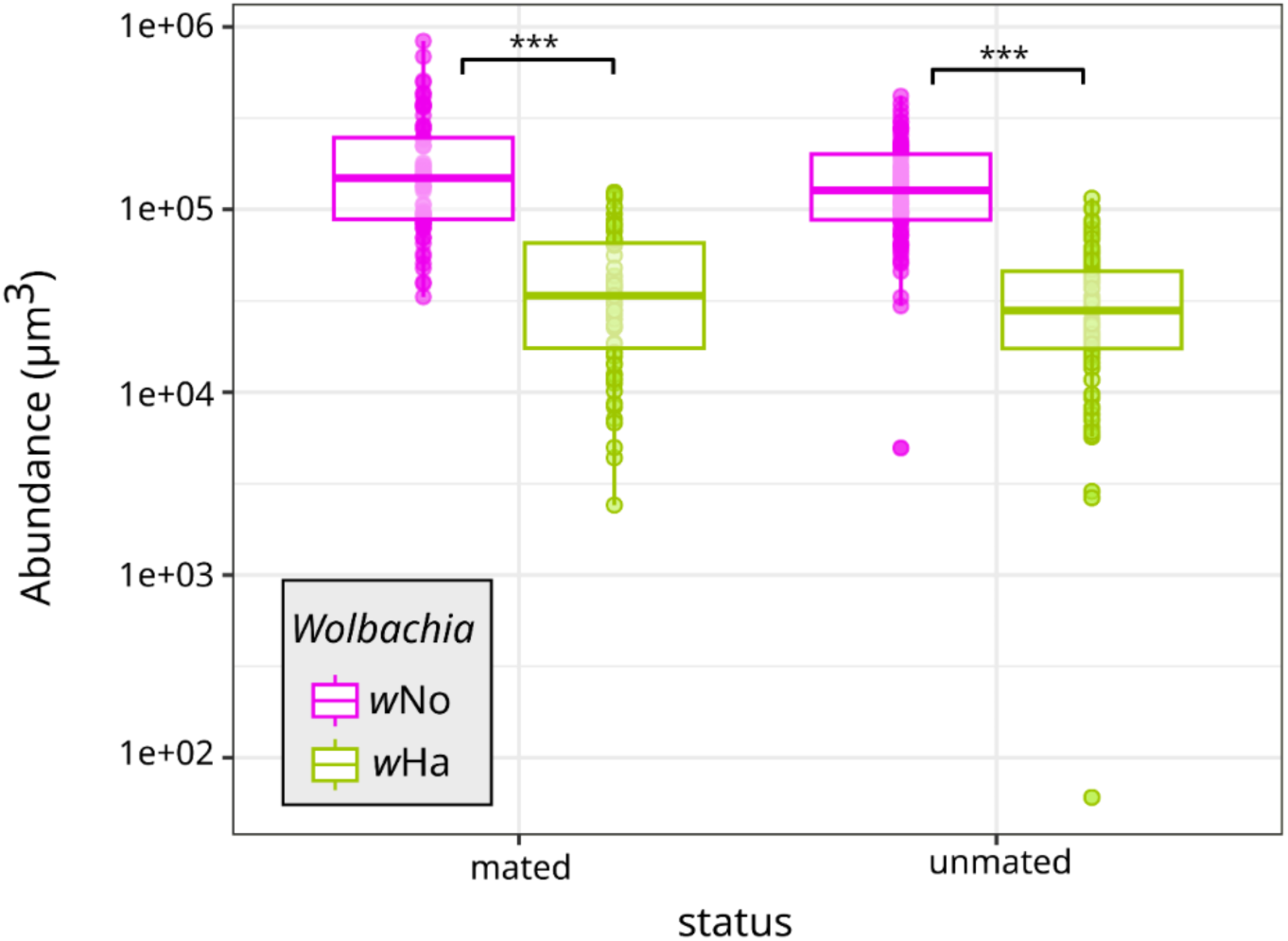
Dynamics of *w*Ha and *w*No in whole ovarioles mirrors the abundance patterns previously observed with qPCR. [48]. To compare strain dynamics observed in qPCR [48] to imaging data, we summed the abundance measurements from each egg chamber within an ovariole to produce a “whole ovariole” abundance measurement. Our FISH-based quantification method found *w*No is approximately four times more abundant than *w*Ha in the ovaries, compared to 2-3 times as abundant when measured with qPCR. Despite differences in quantification method (area occupied by *Wolbachia* vs. number of *Wolbachia* genome copies), we observe that *w*No is significantly more abundant than *w*Ha throughout the ovary. Significance values ***<0.001, **<0.01, *<0.05 calculated from estimated marginal means test with Bonferroni correction for multiple comparisons.

**Supplemental Figure S8.**
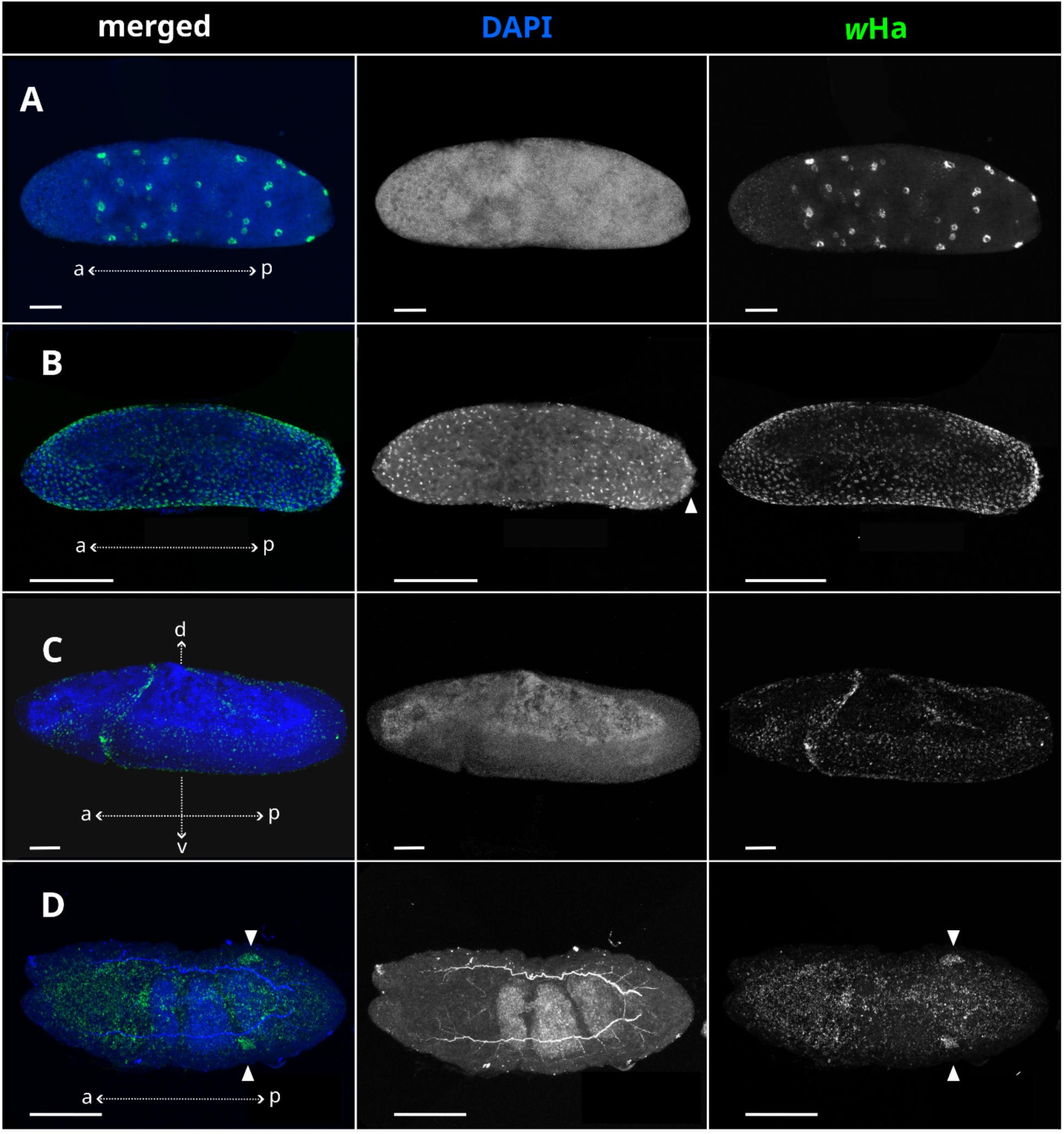
***w*Ha distribution in embryos across development when in monoinfection.** Maximum intensity z-projections of egg chambers, stained with *w*Ha-specific FISH probes and DAPI. (A) Stage 3 embryo, with perinuclear localization of *w*Ha (B) Stage 5 embryo, undergoing formation of the cellular blastoderm and pole cells (arrowhead). (C) Stage 14 embryo, undergoing dorsal closure and with *Wolbachia* localizing to the putative gonadal imaginal discs (arrowheads). (D) Stage 16 embryo, with localization of *w*Ha to the putative gonadal imaginal discs.

## REFERENCES

1. Lindsey AR et al. The intracellular symbiont *Wolbachia* alters *Drosophila* development and metabolism to buffer against nutritional stress. PLOS Genet 2025;21:e1011905. 10.1371/journal.pgen.1011905

2. Douglas AE. Omics and the metabolic function of insect–microbial symbioses. Curr Opin Insect Sci 2018;29:1–6. 10.1016/j.cois.2018.05.012

3. Ankrah NYD et al. Syntrophic splitting of central carbon metabolism in host cells bearing functionally different symbiotic bacteria. ISME J 2020;14:1982–1993. 10.1038/s41396-020-0661-z

4. Brentassi ME, de la Fuente D. Obligate mutualistic heritable symbiosis in sap-feeding insects: an intricate relationship in nature. Symbiosis 2024;92:159–185. 10.1007/s13199-023-00965-1

5. Duron O, Gottlieb Y. Convergence of Nutritional Symbioses in Obligate Blood Feeders. Trends Parasitol 2020;36:816–825. 10.1016/j.pt.2020.07.007

6. Hansen AK, Moran NA. The impact of microbial symbionts on host plant utilization by herbivorous insects. Mol Ecol 2014;23:1473–1496. 10.1111/mec.12421

7. Douglas AE. The microbial dimension in insect nutritional ecology. Funct Ecol 2009;23:38–47. 10.1111/j.1365-2435.2008.01442.x

8. Balmand S et al. Bacterial tubular networks channel carbohydrates in insect endosymbiosis. Cell 2025;188:7355–7365.e16. 10.1016/j.cell.2025.10.001

9. Snyder AK, Rio RVM. “*Wigglesworthia morsitans*” Folate (Vitamin B9) Biosynthesis Contributes to Tsetse Host Fitness. Appl Environ Microbiol 2015;81:5375–5386. 10.1128/AEM.00553-15

10. Fricke LC, Villalta MD, Lindsey AR. Endosymbionts interacting with sex-determining genes and processes. Curr Opin Insect Sci 2025;72:101410. 10.1016/j.cois.2025.101410

11. Perlmutter JI, Bordenstein SR. Microorganisms in the reproductive tissues of arthropods. Nat Rev Microbiol 2020;18:97–111. 10.1038/s41579-019-0309-z

12. Oliver KM et al. Facultative bacterial symbionts in aphids confer resistance to parasitic wasps. Proc Natl Acad Sci 2003;100:1803–1807. 10.1073/pnas.0335320100

13. Lindsey ARI et al. Conflict in the Intracellular Lives of Endosymbionts and Viruses: A Mechanistic Look at *Wolbachia*-Mediated Pathogen-blocking. Viruses 2018;10:141. 10.3390/v10040141

14. Arnam EBV, R. Currie C, Clardy J. Defense contracts: molecular protection in insect-microbe symbioses. Chem Soc Rev 2018;47:1638–1651. 10.1039/C7CS00340D

15. Oliver KM, Smith AH, Russell JA. Defensive symbiosis in the real world – advancing ecological studies of heritable, protective bacteria in aphids and beyond. Funct Ecol 2014;28:341–355. 10.1111/1365-2435.12133

16. Kaltenpoth M, Engl T. Defensive microbial symbionts in Hymenoptera. Funct Ecol 2014;28:315–327. 10.1111/1365-2435.12089

17. Vorburger C, Perlman SJ. The role of defensive symbionts in host–parasite coevolution. Biol Rev 2018;93:1747–1764. 10.1111/brv.12417

18. Holt JR et al. Insect–microbe interactions and their influence on organisms and ecosystems. Ecol Evol 2024;14:e11699. 10.1002/ece3.11699

19. Duron O, Hurst GD. Arthropods and inherited bacteria: from counting the symbionts to understanding how symbionts count. BMC Biol 2013;11:45. 10.1186/1741-7007-11-45

20. Duron O et al. The diversity of reproductive parasites among arthropods: *Wolbachia* do not walk alone. BMC Biol 2008;6:27. 10.1186/1741-7007-6-27

21. Mackevicius-Dubickaja V et al. *Wolbachia* Feminises a Spider Host With Assistance From Co-Infecting Symbionts. Environ Microbiol 2025;27:e70149. 10.1111/1462-2920.70149

22. Moutailler S et al. Co-infection of Ticks: The Rule Rather Than the Exception. PLoS Negl Trop Dis 2016;10:e0004539. 10.1371/journal.pntd.0004539

23. Rio RVM et al. Dynamics of multiple symbiont density regulation during host development: tsetse fly and its microbial flora. Proc R Soc B Biol Sci 2005;273:805–814. 10.1098/rspb.2005.3399

24. Brown AMV et al. Genomic evidence for plant-parasitic nematodes as the earliest *Wolbachia* hosts. Sci Rep 2016;6:34955. 10.1038/srep34955

25. Zug R, Hammerstein P. Still a Host of Hosts for *Wolbachia*: Analysis of Recent Data Suggests That 40% of Terrestrial Arthropod Species Are Infected. PLOS ONE 2012;7:e38544. 10.1371/journal.pone.0038544

26. Kaur R et al. Living in the endosymbiotic world of *Wolbachia*: A centennial review. Cell Host Microbe 2021;29:879–893. 10.1016/j.chom.2021.03.006

27. Ferri E et al. New Insights into the Evolution of *Wolbachia* Infections in Filarial Nematodes Inferred from a Large Range of Screened Species. PLOS ONE 2011;6:e20843. 10.1371/journal.pone.0020843

28. Cordaux R et al. Widespread *Wolbachia* infection in terrestrial isopods and other crustaceans. ZooKeys 2012;123–131. 10.3897/zookeys.176.2284

29. Vancaester E, Blaxter M. Phylogenomic analysis of *Wolbachia* genomes from the Darwin Tree of Life biodiversity genomics project. PLOS Biol 2023;21:e3001972. 10.1371/journal.pbio.3001972

30. Werren JH, Zhang W, Guo LR. Evolution and phylogeny of *Wolbachia*: reproductive parasites of arthropods. Proc R Soc Lond B Biol Sci 1997;261:55–63. 10.1098/rspb.1995.0117

31. Werren JH, Windsor DM. *Wolbachia* infection frequencies in insects: evidence of a global equilibrium? Proc R Soc B Biol Sci 2000;267:1277–1285. 10.1098/rspb.2000.1139

32. Balmand S et al. Tissue distribution and transmission routes for the tsetse fly endosymbionts. J Invertebr Pathol 2013;112:S116–S122. 10.1016/j.jip.2012.04.002

33. Merçot H, Poinsot D. *Wolbachia* transmission in a naturally bi-infected *Drosophila simulans* strain from New-Caledonia. Entomol Exp Appl 1998;86:97–103. 10.1046/j.1570-7458.1998.00269.x

34. Ijichi N et al. Internal Spatiotemporal Population Dynamics of Infection with Three *Wolbachia* Strains in the Adzuki Bean Beetle, *Callosobruchus chinensis* (Coleoptera: Bruchidae). Appl Environ Microbiol 2002;68:4074–4080. 10.1128/AEM.68.8.4074-4080.2002

35. Ant TH, Sinkins SP. A *Wolbachia* triple-strain infection generates self-incompatibility in *Aedes albopictus* and transmission instability in *Aedes aegypti*. Parasit Vectors 2018;11:295. 10.1186/s13071-018-2870-0

36. Lv N et al. Antagonistic interaction between male-killing and cytoplasmic incompatibility induced by *Cardinium* and *Wolbachia* in the whitefly, *Bemisia tabaci*. Insect Sci 2021;28:330–346. 10.1111/1744-7917.12793

37. Takano S et al. Coinfection of *Mesenetia* and rescuing *Wolbachia* in the coconut hispine beetle. Physiol Entomol ;n/a. 10.1111/phen.70003

38. Dobson SL, Rattanadechakul W, Marsland EJ. Fitness advantage and cytoplasmic incompatibility in *Wolbachia* single- and superinfected *Aedes albopictus*. Heredity 2004;93:135–142. 10.1038/sj.hdy.6800458

39. Zhu L-Y et al. *Wolbachia* Strengthens *Cardinium*-Induced Cytoplasmic Incompatibility in the Spider Mite *Tetranychus piercei* McGregor. Curr Microbiol 2012;65:516–523. 10.1007/s00284-012-0190-8

40. James AC et al. Dynamics of double and single *Wolbachia* infections in *Drosophila simulans* from New Caledonia. Heredity 2002;88:182–189. 10.1038/sj.hdy.6800025

41. Mouton L et al. Strain-specific regulation of intracellular *Wolbachia* density in multiply infected insects. Mol Ecol 2003;12:3459–3465. 10.1046/j.1365-294X.2003.02015.x

42. Toomey ME et al. Evolutionarily conserved *Wolbachia*-encoded factors control pattern of stem-cell niche tropism in *Drosophila* ovaries and favor infection. Proc Natl Acad Sci 2013;110:10788–10793. 10.1073/pnas.1301524110

43. Frydman HM et al. Somatic stem cell niche tropism in *Wolbachia*. Nature 2006;441:509–512. 10.1038/nature04756

44. Radousky YA et al. Distinct *Wolbachia* localization patterns in oocytes of diverse host species reveal multiple strategies of maternal transmission. Genetics 2023;224:iyad038. 10.1093/genetics/iyad038

45. Fujiwara A et al. Subcellular Niche Segregation of Co-Obligate Symbionts in Whiteflies. Microbiol Spectr 2022;11:e04684–22. 10.1128/spectrum.04684-22

46. Strunov A et al. Restriction of *Wolbachia* Bacteria in Early Embryogenesis of Neotropical *Drosophila* Species via Endoplasmic Reticulum-Mediated Autophagy. mBio 2022;13:e03863–21. 10.1128/mbio.03863-21

47. Poinsot D, Montchamp-Moreau C, Merçot H. *Wolbachia* segregation rate in *Drosophila simulans* naturally bi-infected cytoplasmic lineages. Heredity 2000;85:191–198. 10.1046/j.1365-2540.2000.00736.x

48. Jones MW et al. Infection Dynamics of Cotransmitted Reproductive Symbionts Are Mediated by Sex, Tissue, and Development. Appl Environ Microbiol 2022;88:e0052922. 10.1128/aem.00529-22

49. Bhattacharya T, Newton ILG, Hardy RW. *Wolbachia* elevates host methyltransferase expression to block an RNA virus early during infection. PLOS Pathog 2017;13:e1006427. 10.1371/journal.ppat.1006427

50. Weisburg WG et al. 16S ribosomal DNA amplification for phylogenetic study. J Bacteriol 1991;173:697–703. 10.1128/jb.173.2.697-703.1991

51. Kuraku S et al. aLeaves facilitates on-demand exploration of metazoan gene family trees on MAFFT sequence alignment server with enhanced interactivity. Nucleic Acids Res 2013;41:W22–W28. 10.1093/nar/gkt389

52. Katoh K, Rozewicki J, Yamada KD. MAFFT online service: multiple sequence alignment, interactive sequence choice and visualization. Brief Bioinform 2019;20:1160–1166. 10.1093/bib/bbx108

53. Nevalainen LB, Layton EM, Newton ILG. *Wolbachia* Promotes Its Own Uptake by Host Cells. Infect Immun 2023;91:e0055722. 10.1128/iai.00557-22

54. Rothwell WF, Sullivan W. Fixation of *Drosophila* Embryos. Cold Spring Harb Protoc 2007;2007:pdb.prot4827. 10.1101/pdb.prot4827

55. Jia D et al. Automatic stage identification of *Drosophila* egg chamber based on DAPI images. Sci Rep 2016;6:18850. 10.1038/srep18850

56. R Core Team. R: A Language and Environment for Statistical Computing. 2024. Vienna, Austria: R Foundation for Statistical Computing, 2024.

57. Lenth R, Piaskowski J. emmeans: Estimated Marginal Means, aka Least-Squares Means. 2026. 2026.

58. Wickham H. ggplot2: elegant graphics for data analysis, Second edition. Cham: Springer international publishing, 2016.

59. Harrington B. Inkscape. 2004. 2004.

60. Zhang C et al. The insect somatostatin pathway gates vitellogenesis progression during reproductive maturation and the post-mating response. Nat Commun 2022;13:969. 10.1038/s41467-022-28592-2

61. Ameku T, Niwa R. Mating-Induced Increase in Germline Stem Cells via the Neuroendocrine System in Female *Drosophila*. PLOS Genet 2016;12:e1006123. 10.1371/journal.pgen.1006123

62. Hoshino R, Niwa R. Regulation of Mating-Induced Increase in Female Germline Stem Cells in the Fruit Fly *Drosophila melanogaster*. Front Physiol 2021;12. 10.3389/fphys.2021.785435

63. Week B et al. Applying evolutionary theory to understand host–microbiome evolution. Nat Ecol Evol 2025;9:1769–1780. 10.1038/s41559-025-02846-w

64. Kittayapong P et al. Combined sterile insect technique and incompatible insect technique: The first proof-of-concept to suppress *Aedes aegypti* vector populations in semi-rural settings in Thailand. PLoS Negl Trop Dis 2019;13:e0007771. 10.1371/journal.pntd.0007771

65. Moreira LA et al. A *Wolbachia* Symbiont in *Aedes aegypti* Limits Infection with Dengue, Chikungunya, and *Plasmodium*. Cell 2009;139:1268–1278. 10.1016/j.cell.2009.11.042

66. Hoffmann AA et al. Successful establishment of *Wolbachia* in *Aedes* populations to suppress dengue transmission. Nature 2011;476:454–457. 10.1038/nature10356

67. McCall K. Eggs over easy: cell death in the *Drosophila* ovary. Dev Biol 2004;274:3–14. 10.1016/j.ydbio.2004.07.017

68. Riechmann V, Ephrussi A. Axis formation during *Drosophila* oogenesis. Curr Opin Genet Dev 2001;11:374–383. 10.1016/S0959-437X(00)00207-0

69. Weil TT et al. Changes in bicoid mRNA Anchoring Highlight Conserved Mechanisms during the Oocyte-to-Embryo Transition. Curr Biol 2008;18:1055–1061. 10.1016/j.cub.2008.06.046

70. Warrior R. Primordial Germ Cell Migration and the Assembly of the *Drosophila* Embryonic Gonad. Dev Biol 1994;166:180–194. 10.1006/dbio.1994.1306

71. Veneti Z et al. Heads or Tails: Host-Parasite Interactions in the *Drosophila*-*Wolbachia* System. Appl Environ Microbiol 2004;70:5366–5372. 10.1128/AEM.70.9.5366-5372.2004

72. Rice DW, Sheehan KB, Newton ILG. Large-Scale Identification of *Wolbachia pipientis* Effectors. Genome Biol Evol 2017;9:1925–1937. 10.1093/gbe/evx139

73. Mercot H et al. Variability within the Seychelles Cytoplasmic Incompatibility System in *Drosophila Simulans*. Genetics 1995;141:1015–1023. 10.1093/genetics/141.3.1015

